# An ER phospholipid hydrolase drives ER-associated mitochondrial constriction for fission and fusion

**DOI:** 10.1101/2022.10.24.513548

**Authors:** Tricia T. Nguyen, Gia K. Voeltz

## Abstract

Mitochondria are dynamic organelles that undergo cycles of fission and fusion at a unified platform defined by endoplasmic reticulum (ER)-mitochondria membrane contact sites (MCSs). These MCSs or nodes co-localize fission and fusion machinery. We set out to identify how ER-associated mitochondrial nodes can regulate both fission and fusion machinery assembly. We have used a promiscuous biotin ligase linked to the fusion machinery, Mfn1, and proteomics to identify an ER membrane protein, Aphyd, as a major regulator of node formation. In the absence of Aphyd, fission and fusion machineries fail to recruit to ER-associated mitochondrial nodes and fission and fusion rates are significantly reduced. Aphyd contains an acyltransferase motif and an α/β hydrolase domain and point mutations in critical residues of these regions fail to rescue the formation of ER-associated mitochondrial hot spots. These data suggest a mechanism whereby Aphyd functions by altering phospholipid composition at ER-mitochondria MCSs. Our data present the first example of an ER membrane protein that regulates the recruitment of both fission and fusion machineries to mitochondria.

## Introduction

Cells maintain a characteristic mitochondrial architecture important for cellular metabolism and function. Mitochondria maintain their overall architecture and morphology by undergoing cycles of fission and fusion (Friedman et al., 2010; Twig et al., 2008; Youle and Van Der Bliek, 2012). Disruption of these cycles results in fragmentation or elongation, which can be detrimental to cell health and is associated with various disease states (Rambold et al., 2011; Wai and Langer, 2016). Both processes are first initiated by the endoplasmic reticulum (ER) at ER-mitochondria membrane contact sites (MCSs). Several factors have been linked to ER-associated mitochondrial fission, including two actin nucleators, INF2 and Spire1c, which are proposed to polymerize actin at ER-mitochondria MCSs to initiate constriction of the outer mitochondrial membrane (OMM) (Korobova et al., 2013; Manor et al., 2015). Subsequently, two GTPases (Drp1 and Dyn2) are recruited to the OMM at ER MCSs to further constrict the OMM, which leads to mitochondrial division (Bleazard et al., 1999; Ferguson and De Camilli, 2012; Labrousse et al., 1999; Lee et al., 2016; Smirnova et al., 1998). Several factors have also been linked to ER-associated mitochondrial fusion. ER-mitochondria MCSs first dictate sites where OMM fusion occurs via Mfn1/2 oligomerization in trans to drive membrane fusion upon GTP hydrolysis (Abrisch et al., 2020; Chen et al., 2003; Guo et al., 2018; Santel and Fuller, 2001). Subsequently, inner mitochondrial membrane (IMM) fusion occurs via the GTPase, Opa1 (Ban et al., 2017; Herlan et al., 2003; Lee et al., 2004; Legros et al., 2002; Meeusen et al., 2006; Misaka et al., 2002). However, how the ER contributes to defining a site on mitochondria that is primed for constriction and sufficient to coordinate the recruitment of both fission and fusion machineries and further how these two seemingly opposing activities are co-recruited and also balanced is unclear.

We have recently discovered that cycles of fission and fusion occur at the same location or hot spots that are spatially dictated by the ER. These predefined branch points, or nodes, are where both fission (Drp1) and fusion (Mfn1) machineries converge (Abrisch et al., 2020). However, it is not known how these nodes are formed or regulated. We hypothesized that an ER membrane protein facilitates the formation of these nodes. To identify an ER machinery involved in node formation, we have taken advantage of a promiscuous biotin ligase (TurboID) that can biotinylate proteins within a ~10-30 nm range upon biotin addition (Branon et al., 2018; Roux et al., 2012). We have fused TurboID to the known fusion machinery, Mfn1, to biotinylate and subsequently identify neighboring ER proteins that could regulate these nodes. Using this strategy, we have identified an ER membrane-localized lipid hydrolase and acyltransferase, Aphyd, that alters lipid membranes for mitochondrial constriction after ER contact sites are established. Aphyd is not only required for ER-associated mitochondrial constriction but it is also the first ER protein to be shown to be required for both fission and fusion machinery to assemble at contact sites.

## Results

### Identification of ER-localized lipid hydrolase, Aphyd, by proximal proteomics

Membrane contact sites (MCSs) between the ER and mitochondria define the position where mitochondria undergo fission and fusion (Abrisch et al., 2020; Friedman et al., 2011; Guo et al., 2018). Surprisingly, the fission and fusion machineries co-localize at the same ER-mitochondria MCSs, not separate ones (Abrisch et al., 2020). It is not known how the ER functions mechanistically to define constriction sites on mitochondria for both fission and fusion machinery recruitment. Here, we sought to identify ER factors that regulate the assembly of ER-associated nodes where both mitochondrial fission and fusion machinery are recruited. Our strategy was to fuse TurboID, an optimized promiscuous biotin ligase, to the outer mitochondrial membrane (OMM) fusion protein, Mfn1 (Figure 1A), which assembles at ER-mitochondria MCSs (Abrisch et al., 2020; Branon et al., 2018). TurboID can biotinylate proteins within a ~10-30 nanometer range upon addition of biotin (Branon et al., 2018; Roux et al., 2012). By immunofluorescence, V5-TurboID-Mfn1 (magenta, with V5 antibody) co-localized well with GFP-Mfn1 (green) puncta on mitochondria (mito-BFP, blue) in U-2 OS cells (Figure 1B, top panels). As a negative control, we similarly fused TurboID to a GTPase domain mutant Mfn1E209A (Figure 1A and 1B). This mutant is deficient in its ability to hydrolyze GTP, does not homodimerize, cannot drive fusion (Cao et al., 2017; Sloat et al., 2019), and does not enrich at ER-mitochondria MCSs with WT GFP-Mfn1 puncta (compare magenta to green, Figure 1B, bottom panels) consistent with previous reports (Abrisch et al., 2020).

**Figure 1.**
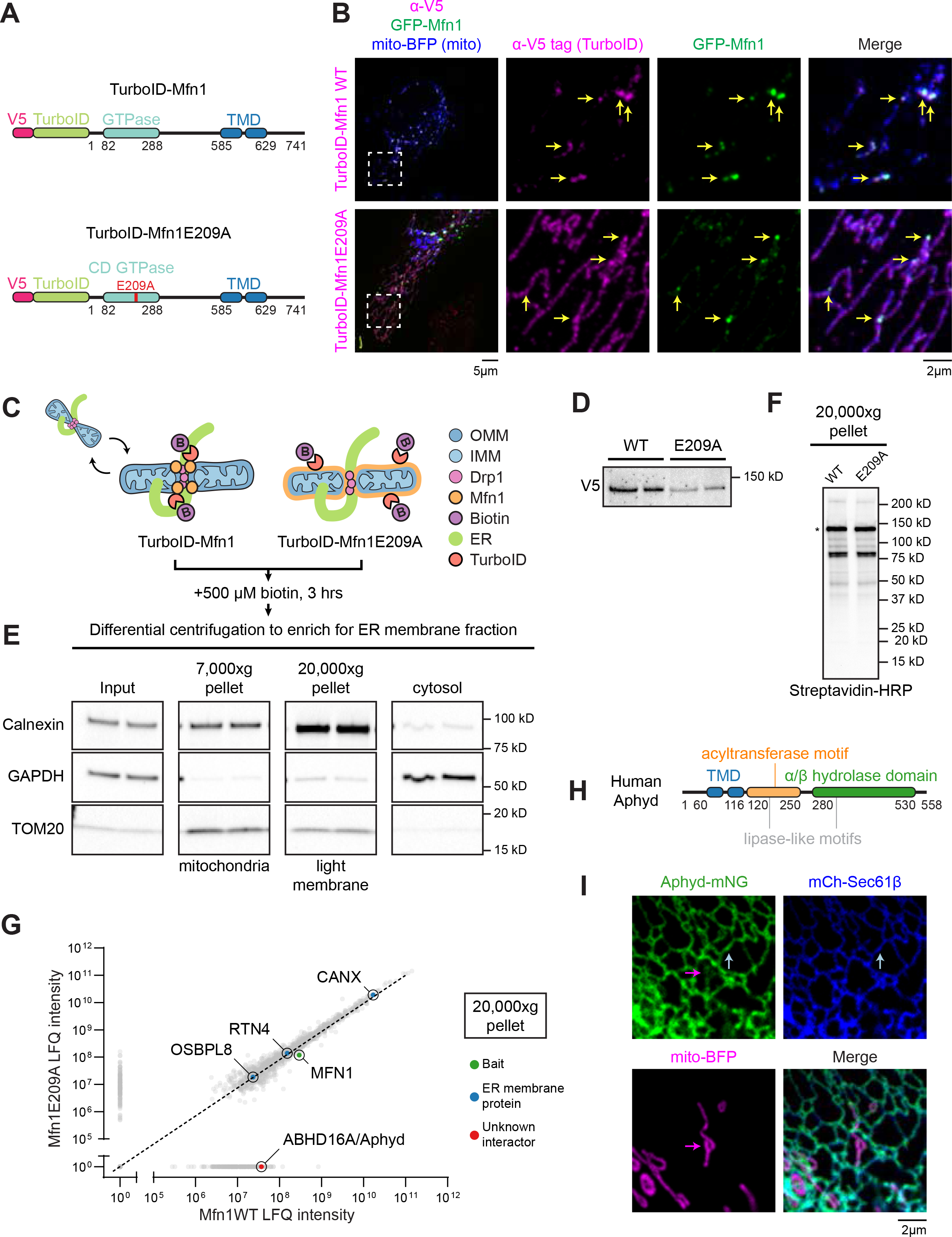
Identification of ER-localized lipid hydrolase, Aphyd, by proximal proteomics. (**A**) Cartoon diagram and domain organization of V5-TurboID-Mfn1 and V5-TurboID-Mfn1E209A (human Mfn1 and indicated amino acid numbers). V5-tagged TurboID was added to the N-terminus of Mfn1. Red E209A indicates the catalytically dead (CD) mutation created in the GTPase domain. TMD: Transmembrane domain. (**B**) Representative images and insets of a HeLa cell expressing GFP-Mfn1 (green), mito-BFP (blue), and immunostained with antibody against the V5 tag (magenta, to detect Mfn1 constructs). Yellow arrows indicate GFP-Mfn1 puncta along mitochondria. (**C**) Cartoon diagram of strategy used in HeLa cells to biotinylate ER-mitochondria MCS proteins with V5-TurboID-Mfn1 vs. V5-TurboID-Mfn1E209A. OMM: outer mitochondrial membrane (blue); IMM: inner mitochondrial membrane (light blue); Drp1: Dynamin-related protein 1 (fission machinery, pink); Mfn1: Mitofusin 1 (fusion machinery, orange); B: biotin; ER: endoplasmic reticulum. (**D**) Immuno-blot analysis (anti-V5) shows relative expression of V5-TurboID-Mfn1 and V5-TurboID-Mfn1E209A in HeLa cells. (**E**) Immuno-blot analyses of fractions collected by differential centrifugation including: 7,000 x g pellet containing mitochondria (TOM20, mitochondria), 20,000 x g pellet containing light membrane and ER (Calnexin, ER), and supernatant containing cytosol (GAPDH, cytosol). (**F**) The 20,000 x g pellet was solubilized and probed with Streptavidin HRP to reveal biotinylation profiles for each sample prior to mass spectrometry. Asterisk denotes a band size indicative of construct self biotinylation. (**G**) Average LFQ intensities of proteins biotinylated and enriched in Mfn1 WT vs. Mfn1E209A sample in the 20,000 x g pellet. Dashed line indicates equivalent enrichment in the WT vs. E209A mutant sample. Data from LFQ intensity is the average of two technical replicates. CANX: Calnexin; RTN4: Reticulon-4; MFN1: Mitofusin-1; OSBPL8: Oxysterol binding protein like 8; Aphyd: Alpha/beta phospholipid hydrolase (previously named ABHD16A, Alpha/beta hydrolase domain containing phospholipase 16A). (**H**) Cartoon diagram of human Aphyd with motif or domain annotations. (**I**) Representative inset of a U-2 OS cell expressing low levels of Aphyd-mNG (green), mCh-Sec61β (ER, blue), and mito-BFP (mitochondria, magenta). Arrows indicate Aphyd localization to ER (blue) and mitochondria (magenta). Scale bar = 5 μm or 2 μm for insets. See Figure 1-source data 1-10.

TurboID proximity biotinylation experiments were performed by transfecting HeLa cells with either V5-TurboID-Mfn1 or V5-TurboID-Mfn1E209A followed by treatment with 500 μM biotin for 3 hours (Figure 1C). After biotinylation, the ER membrane fraction was enriched by differential centrifugation as described previously (Hoyer et al., 2018; Wieckowski et al., 2009). Light membranes were pelleted at 20,000 x g and immunoblot analysis was used to confirm that both fractions were enriched with ER and mostly depleted of other organelles and cytosol (Figure 1D and 1E). The biotinylation profile of the light membrane fraction with a streptavidin-HRP probe confirmed that both fusion constructs were enzymatically active (Figure 1F). The resulting biotinylated light membrane fraction was purified on streptavidin columns and analyzed by mass spectrometry. Similar self-biotinylation levels of Mfn1 WT (wild type) versus mutant were seen in each condition via LFQ intensity (Figure 1G). The ER membrane fraction was equivalently enriched in both conditions, because known ER membrane proteins, such as calnexin (CANX), Reticulon-4 (RTN4), and Oxysterol binding protein like 8 (OSBPL8/ORP8), were identified to equal levels (Figure 1G). Additionally, we identified a novel ER membrane protein, ABHD16A, (renamed here Aphyd for <u>A</u>lpha/beta <u>p</u>hospholipid <u>hy</u>drolase for simplicity) that was highly enriched in the WT Mfn1 sample (19^th^ highest enrichment) as a candidate effector of ER-associated mitochondrial node formation (Figure 1G).

Aphyd is reported to be a phospholipid hydrolase that resides on the ER membrane via two transmembrane domains (Lord et al., 2013; Singh et al., 2020). It is highly conserved in vertebrates (96% conserved from human to mouse) and contains an alpha/beta hydrolase domain responsible for its phospholipid hydrolase activity, a predicted acyltransferase motif, and two predicted lipase-like motifs (Figure 1H; Kamat et al., 2015; Lord et al., 2013; Singh et al., 2020). Phospholipid hydrolases are known to remove one acyl chain from phospholipids, whereas, acyltransferase motifs catalyze the opposite reaction of a phospholipid hydrolase: where a single chain phospholipid reacts with an acyl-CoA (coenzyme A) to restore a dual chain phospholipid (Aguado and Campbell, 1998; Eberhardt et al., 1997; Lord et al., 2013; Zhao et al., 2008).

We generated a fluorescently tagged Aphyd (Aphyd-mNeonGreen) to determine its localization in U-2 OS cells by live confocal fluorescence microscopy. U-2 OS cells were co-transfected with low levels of Aphyd-mNeonGreen (green), a mitochondrial matrix marker (mito-BFP, magenta), and an ER membrane marker (mCherry-Sec61β, blue). At low expression levels, Aphyd-mNG co-localized homogenously along the ER membrane (blue arrow) and to a much lesser degree on mitochondria (magenta arrow), but did not appear to accumulate at MCSs (Figure 1I and S1A). Endogenous Aphyd localization was also analyzed by immunoblot analysis of pure, crude, and mitochondrial-associated membrane (MAM) fractions isolated from U-2 OS cells. These data showed that endogenous Aphyd can be found in the MAM similar to other ER proteins and to a lesser extent in the pure mitochondrial fraction (Figure S1B) (Wieckowski et al., 2009).

### Aphyd is required for formation of ER-associated fission and fusion hot spots

Aphyd was preferentially biotinylated by TurboID-Mfn1 WT. Therefore, we assayed whether Aphyd depletion affects the recruitment of the OMM fusion machinery, Mfn1, to ER-mitochondria MCSs. U-2 OS cells were co-transfected with fluorescently tagged Mfn1 (GFP-Mfn1, green), a mitochondrial matrix marker (mito-BFP, magenta), and an ER marker (mNG-Sec61β, blue) and with either control (siCTRL) or Aphyd siRNA. Immunoblot analysis reveals that Aphyd can be efficiently depleted by siRNA transfected into U-2 OS cells (Figure S2D). Mfn1 recruitment efficiency was scored as the number of Mfn1 puncta present per micron of mitochondrial length. In siCTRL-treated cells, GFP-Mfn1 accumulates at puncta along mitochondria (Figure 2A, top panel), as expected (Abrisch et al., 2020). Aphyd depletion significantly reduced Mfn1 puncta compared to siCTRL-treated cells (Figure 2A and 2D: ~0.1 versus 0.3 puncta per μm of mitochondria, respectively). Mfn1 puncta were restored by re-expression of an siRNA-resistant Aphyd-Halo construct (Figure 2A, 2D, and S2A). These data demonstrate that Aphyd regulates localization of Mfn1 to ER-mitochondria contact sites.

**Figure 2.**
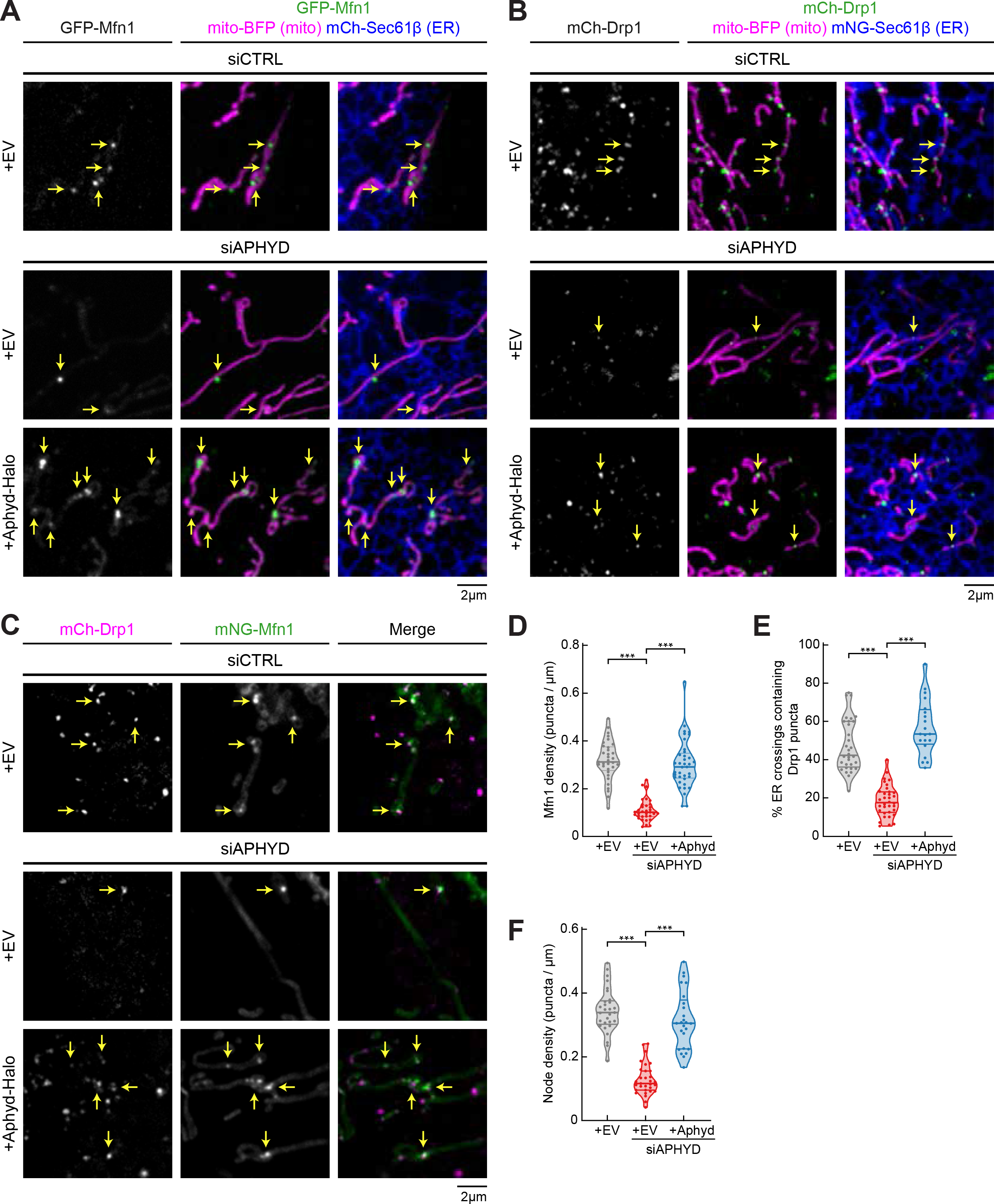
Aphyd is required for formation of ER-associated fission and fusion hot spots. (**A**) Representative images of U-2 OS cells transfected with GFP-Mfn1 (green), mito-BFP (magenta), mCh-Sec61β (ER, blue), and either control siRNA (n=33 cells, top), Aphyd siRNA (n=33 cells, middle), or Aphyd siRNA rescued with Aphyd-Halo (n=37 cells, bottom). Yellow arrows indicate examples of Mfn1 puncta along mitochondria at ER-mitochondria crossings. (**B**) Representative images of U-2 OS cells transfected with mCh-Drp1 (green), mito-BFP (magenta), mNG-Sec61β (ER, blue), and either control siRNA (n=32 cells, top), Aphyd siRNA (n=34 cells, middle), or Aphyd siRNA rescued with Aphyd-Halo (n=20 cells, bottom). Yellow arrows indicate examples of Drp1 puncta at ER-mitochondria crossings. (**C**) Representative images of U-2 OS cells transfected with mCh-Drp1 (magenta), GFP-Mfn1 (green), mito-BFP (not shown), and either control siRNA (n=29 cells, top), Aphyd siRNA (n=31 cells, middle), or Aphyd siRNA rescued with Aphyd-Halo (n=25 cells, bottom). Yellow arrows indicate examples of nodes along mitochondria. (**D**) Quantification of Mfn1 density along mitochondrial length from experiments shown in (A). (**E**) Quantification of percent ER crossings containing Drp1 puncta from experiments shown in (B). (**F**) Quantification of node density along mitochondrial length from experiments shown in (C). All data taken from 3 biological replicates; statistical significance calculated by one-way ANOVA. ***p≤0.001. Scale bar = 2 μm. See Figure 2-source data 1.

Since fission machinery has been shown to co-localize with fusion machinery, we tested whether Aphyd depletion similarly disrupts the recruitment of fission machinery, Drp1, to ER-mitochondria MCSs. U-2 OS cells were co-transfected with fluorescently tagged Drp1 (mCh-Drp1, green), mito-BFP (magenta), and mNG-Sec61β (ER, blue) and with either siCTRL or Aphyd siRNA (Figure 2B and S2B). The efficiency of fission machinery recruitment was scored as the percent of ER-mitochondria crossings that co-localize with a Drp1 puncta. Drp1 recruitment to crossings was significantly reduced in Aphyd-depleted versus siCTRL-treated cells (18% versus 47%, respectively, Figure 2E). The recruitment of Drp1 to ER-associated puncta could be restored by reintroduction of an siRNA-resistant Aphyd expression construct (Aphyd-Halo) (Figure 2B, 2E, and S2B). These data reveal that Aphyd is also required for Drp1 fission machinery recruitment to ER-mitochondria MCSs.

Next, we measured whether Aphyd is required for the assembly of ER-associated mitochondrial nodes, which are locations where Drp1 and Mfn1 machineries co-localize (Abrisch et al., 2020). Cells were co-transfected with mCh-Drp1 (magenta), mNG-Mfn1 (green), and mito-BFP (not shown) and with either siCTRL or Aphyd siRNA and imaged live (Figure 2C and S2C). Node density was scored as the number of puncta containing both Drp1 and Mfn1 per μm of mitochondrial length. Aphyd-depleted cells had significantly fewer nodes than control cells (~0.13 vs. ~0.34 puncta per μm of mitochondria, Figure 2F). Mitochondrial node formation could be rescued by re-expression of siRNA-resistant Aphyd-Halo but not the empty vector (EV) control (Figure 2C, 2F, and S2C). Together, these data demonstrate that Aphyd is required for the co-localization of fission and fusion machineries to mitochondrial nodes, suggesting a common mechanism is required for their recruitment.

### Aphyd is required for efficient cycles of ER-mediated mitochondrial fission and fusion

We predicted that Aphyd depletion would also reduce mitochondrial fission and fusion rates concomitant with its deleterious effect on fission and fusion machinery recruitment to ER-associated nodes. To test this directly, we first scored fusion rates by using a photoconvertible fluorophore to label the outer mitochondrial membrane (mMAPLE-OMP25, as previously described(Abrisch et al., 2020). Briefly, upon 405 nm laser stimulation, mMAPLE fluoresces red instead of green and fusion can be scored by red fluorescence diffusion (Figure S3A). Cells were co-transfected with mMAPLE-OMP25 and either siCTRL or Aphyd siRNA, single mitochondria were photoconverted from green to red (magenta in panels), and fusion rates were scored visually and quantitatively during live 5-minute time-lapse movies by the observation of fluorescence mixing between two mitochondria (Figure 3A-D and S3B). Fusion rates were significantly reduced in Aphyd-depleted cells (Figure 3A, 3B, 3D, and S3B: from a rate of 0.25 (siCTRL) to 0.06 (siAPHYD) fusion events per mitochondrion/minute). Fusion rates could be rescued by re-introduction of siRNA-resistant Aphyd-Halo to depleted cells (Figure 3C, 3D and S3B).

**Figure 3.**
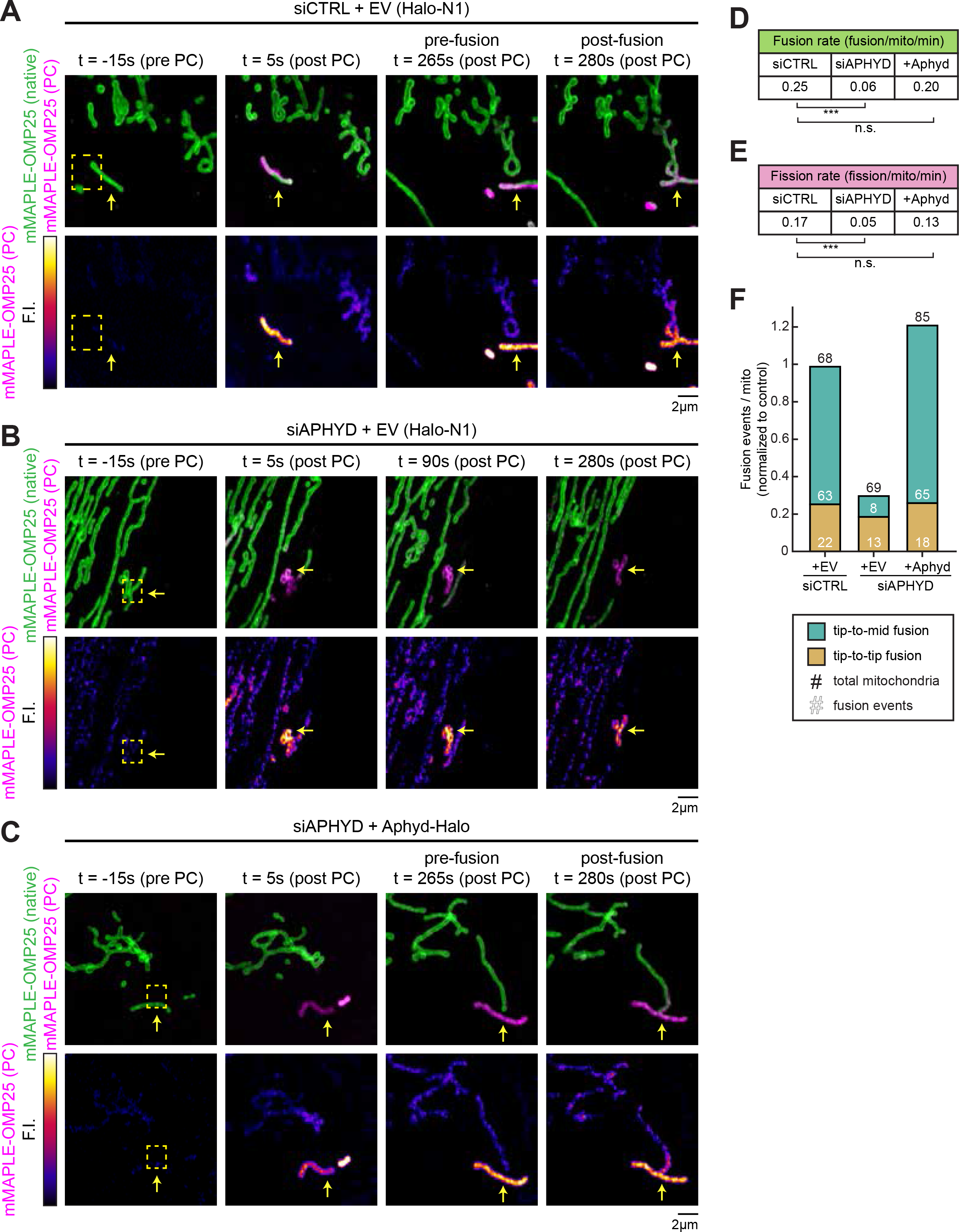
Aphyd is required for efficient cycles of ER-mediated mitochondrial fission and fusion. (**A**) Representative time lapse of a live U-2 OS cell over a 5-minute movie expressing mMAPLE-OMP25 and control siRNA (n=30 cells). Top panel: Green shows native mMAPLE-OMP25 signal while magenta shows photoconverted (PC) mMAPLE-OMP25. Bottom panel: Fire lookup table (LUT) of PC mMAPLE-OMP25 to show fusion event. Yellow box indicates ROI exposed to 405 nm light. Yellow arrow indicates photoconverted mitochondrion. (**B**) As in (A) for live U-2 OS cell expressing mMAPLE-OMP25 and Aphyd siRNA (n=29 cells). Fire LUT and yellow arrow show no fusion occurring over the 5-minute movie. (**C**) As in (A) for live U-2 OS cell expressing mMAPLE-OMP25, Aphyd siRNA, and rescued with Aphyd-Halo (n=31 cells). (**D**) Quantification of fusion rate per mitochondrion per minute from experiments shown in (A-C). (**E**) Quantification of fission rate per mitochondrion per minute from experiments shown in (A-C). (**F**) Quantification of types of fusion events per mitochondrion normalized to siCTRL from experiments shown in (A-D). All data taken from 3 biological replicates; statistical significance calculated by one-way ANOVA. n.s., not significant; ***p≤0.001. Scale bar = 2 μm. See Figure 3-source data 1.

To further investigate the mechanism by which Aphyd depletion disrupts mitochondrial fusion, we binned mitochondrial fusion events into two categories: tip-to-middle fusion and tip-to-tip fusion. Tip-to-middle fusion occurs when one mitochondrial tip marked with Mfn1 travels to a Mfn1-marked spot along the middle of a mitochondrion. Tip-to-tip fusion occurs when two mitochondrial tips marked with Mfn1 come together to fuse. In siCTRL cells, the majority of events occur by tip-to-mid fusion (74%, Figure 3F). Upon Aphyd depletion, tip-to-middle fusion events are highly reduced (63 events vs. 8 events; an 87% reduction) whereas tip-to-tip fusion is less affected (22 events vs. 13 events, a 40% reduction, Figure 3F). What makes these results notable is that the “middle” part of the tip-to-middle fusion event occur at ER-associated mitochondrial constriction sites where fission and fusion machineries also assemble and co-localize (Abrisch et al., 2020).

Next, we scored the effect of Aphyd depletion on the rate of fission (within the same experiment that was used to score fusion rates). Indeed, fission rates were also significantly reduced upon Aphyd depletion (an >3-fold reduction from 0.17 to 0.05 fission events per mitochondrion/minute) (Figure 3E and S3C). Fission rates could also be restored by expressing exogenous siRNA-resistant Aphyd-Halo and not with an EV control (Halo-N1) (Figure 3E and S3C). These data show that Aphyd depletion reduces fusion and fission rates concomitant with the reduced recruitment of fusion and fission machineries to ER-associated nodes. Taken together, Aphyd is a regulator of ER-associated mitochondrial fission and fusion dynamics.

### ER-localized Aphyd is required to maintain mitochondrial morphology

Since Aphyd depletion alters fission and fusion machinery recruitment, we scored the overall effect of Aphyd depletion on mitochondrial morphology as previously described (Lee et al., 2016). U2-OS cells were co-transfected with mito-BFP and either siCTRL or Aphyd siRNA to deplete Aphyd. On average, Aphyd-depleted cells had highly elongated mitochondria compared to control (4.4 μm^2^ versus 2.2 μm^2^, respectively, Figure 4B, 4D, 4E, and S4A). Morphology could be restored to siCTRL lengths by re-expressing siRNA-resistant Aphyd (Aphyd-mNG) (Figure 4C and 4D; also see immunoblot in Figure 4E for relative expression levels). The depletion of Aphyd also caused mitochondrial elongation (compared to control) in HeLa cells, showing that this effect is not cell-type specific (Figure S4C-S4E). Since Aphyd depletion significantly impairs tip-to-middle fusion and overall fission rates, we reasoned that the residual tip-to-tip mitochondrial fusion (which can still occur upon depletion) is sufficient to promote the elongated mitochondrial phenotype.

**Figure 4.**
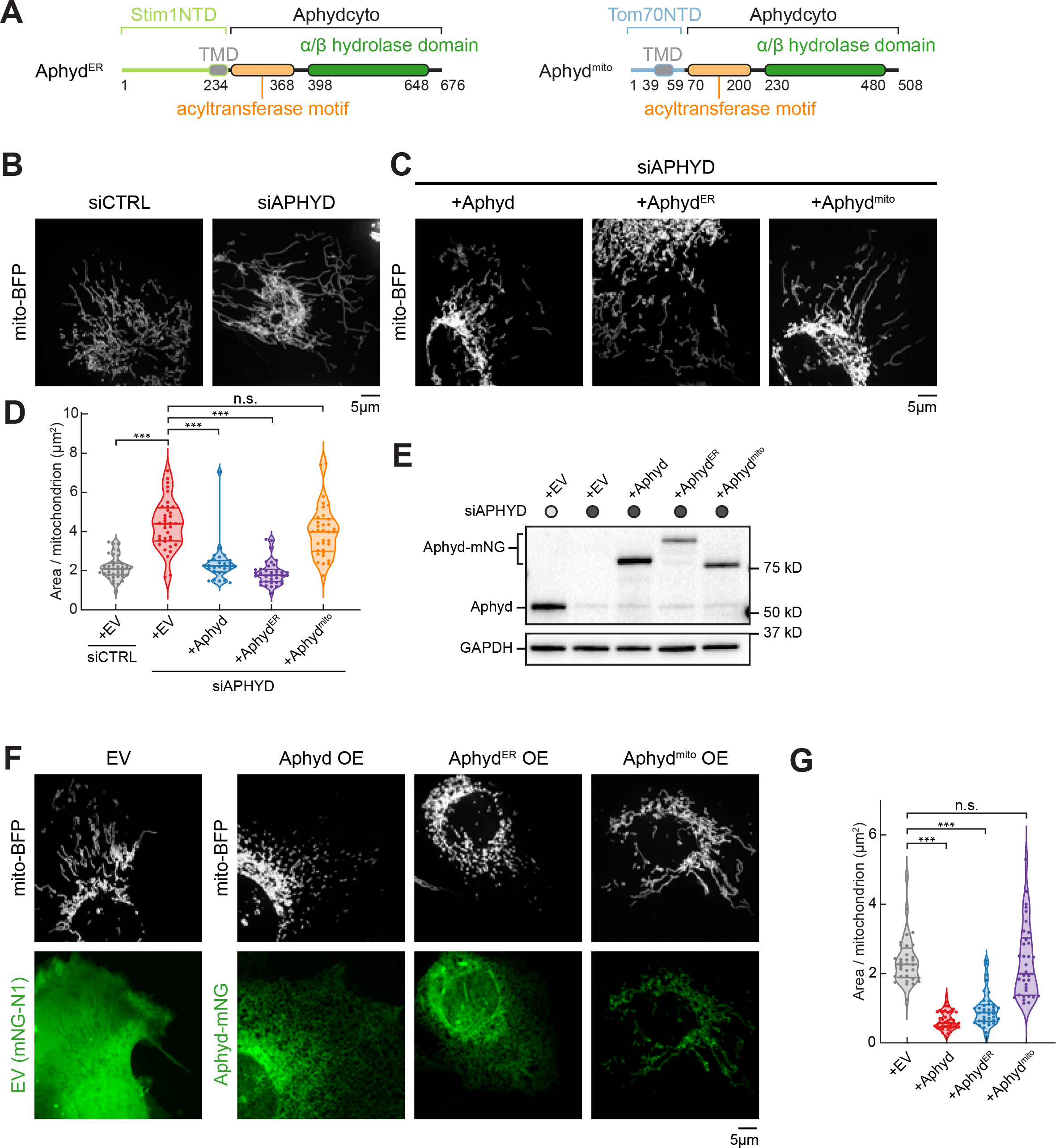
ER-localized Aphyd is required to maintain mitochondrial morphology. (**A**) Domain organization of chimeric constructs for ER-localized Aphyd (Aphyd^ER^) via Stim1’s N-terminal domain (NTD) (left) or mitochondrial-localized Aphyd (Aphyd^mito^) via Tom70’s NTD (right). TMD: Transmembrane domain. (**B**) Representative images of mitochondrial morphology (labeled by mito-BFP, grey) of U-2 OS cells transfected with control siRNA (siCTRL, n=41 cells, left) or Aphyd siRNA (siAPHYD, n=36 cells, right). (**C**) Representative images of mitochondrial morphology (labeled by mito-BFP, grey) of U-2 OS cells transfected with Aphyd siRNA and rescued with either siRNA-resistant Aphyd-mNG (n=28 cells, left), Aphyd^ER^-mNG (n=36 cells, middle), or Aphyd^mito^-mNG (n=34 cells, right). (**D**) Quantification of mean mitochondrial size (area per mitochondrion in μm^2^) within a 15×15 μm ROI from (B) and (C). (**E**) Immunoblot shows efficiency of depletion in cells treated with control siRNA or Aphyd siRNA and rescued with siRNA-resistant Aphyd-mNG constructs. GAPDH serves as a loading control. (**F**) Representative images of mitochondrial morphology of U-2 OS cells (labeled by mito-BFP, grey) and either empty vector (EV, control, n=32 cells), Aphyd-mNG (n=40 cells), Aphyd^ER^-mNG (n=38 cells), or Aphyd^mito^-mNG (n=34 cells) in green. (**G**) Quantification of mean mitochondrial size (area per mitochondrion in μm^2^) within a 15×15 μm ROI from (F). All data taken from 3 biological replicates; statistical significance calculated by one-way ANOVA. n.s., not significant; ***p≤0.001. Scale bar = 5 μm. See Figure 4-source data 1-4.

Our localization experiments revealed that although Aphyd predominantly localizes to the ER, a small amount of Aphyd can also be seen on mitochondria (Fig 1I, S1A, and S1B). To understand how Aphyd functions mechanistically, it is thus important to understand whether the ER-localized or the trace mitochondrial-localized Aphyd pool is responsible for regulating mitochondrial dynamics. We therefore tested whether expression of an exclusively ER-targeted or mitochondrial-targeted Aphyd functions to rescue mitochondrial morphology. To target Aphyd exclusively to the ER or to mitochondria, we replaced the N-terminal transmembrane domains (TMDs) of Aphyd with either the N-terminal TMD of the ER-localized protein Stim1 or the N-terminal TMD of the mitochondrial-localized protein Tom70 (Figure 4A). Both chimeric constructs localize as expected exclusively to the ER or mitochondria, respectively (Figure S4B). However, only re-expression of the siRNA-resistant ER-localized Aphyd (Aphyd^ER^) rescues mitochondrial morphology, while rescue with the mitochondria-localized version (Aphyd^mito^) does not (Figure 4C-4E). In a complementary experiment, we also scored the effect of Aphyd overexpression (OE) on mitochondrial morphology. On average, Aphyd OE cells had a 3.6-fold reduction in area per mitochondrion (Figure 4F and 4G). Consistent with rescue data above (Figure 4B-E, S4A, and S4B), only the ER-localized Aphyd (Aphyd^ER^) drove mitochondrial fragmentation, whereas the mitochondrial-localized form (Aphyd^mito^) did not (Figure 4F and 4G). Together, the depletion and overexpression experiments suggest that the ER-localized Aphyd regulates mitochondrial morphology from the ER.

### Aphyd is not a tether

Our data demonstrate that Aphyd is required for fission and fusion machinery recruitment to ER-mitochondria MCSs. We tested whether this is because Aphyd is required for tethering between the ER and mitochondria. Aphyd was depleted by siRNA and a dimerization-dependent fluorescent protein (ddFP) reporter system was used to score ER-mitochondria MCS, as previously described (Abrisch et al., 2020). This reporter system consists of a heterodimeric fluorescent protein complex (dimly red fluorescent RA and protein binding partner B) (Ding et al., 2015). Upon dimerization of RA with B, RA fluorescence increases leading to a detectable red signal. The low K_d_ of the RA/B interaction (~7 μM) ensures that the interaction is reversible and does not cause artificial wrapping, which allows for bona-fide MCSs to be visualized. RA was fused to an ER membrane protein, Sec61β, to target half of the dimer to the ER and B was fused to the OMM protein Mff, as previously described (Abrisch et al., 2020). When the ER and mitochondria come into close proximity (tethering distance), an increased fluorescence intensity is observed (Figure 5A). Cells were co-transfected with either control or Aphyd siRNA, mito-BFP (magenta), an ER marker (mNG-KDEL, blue), ddFP reporters (RA-Sec61β and B-Mff, dimer will fluoresce green), and were rescued with siRNA-resistant Aphyd (Aphyd-SNAP) or an EV control (SNAP-N1) (Figure 5B, S5A, and S5B). ER-mitochondria contact was quantified by assessing binary ddFP overlap with mitochondria as a proportion of total mitochondrial area within a 20×20 μm^2^ region of interest (ROI). To ensure resolvable regions were quantified in each condition, we also measured the binary ER overlap with mitochondria as a proportion of total mitochondrial area, which was notably similar throughout each condition (Figure 5C). Thus, Aphyd depletion leads to 2.1-fold increase in ER-mitochondria MCS area (ddFP) compared to siCTRL-treated cells (Figure 5D). This increased contact was reduced by re-expression of siRNA-resistant Aphyd (Figure 5B, 5D, S6A, and S6B). These data suggest that Aphyd is not an ER-mitochondria contact site tether.

**Figure 5.**
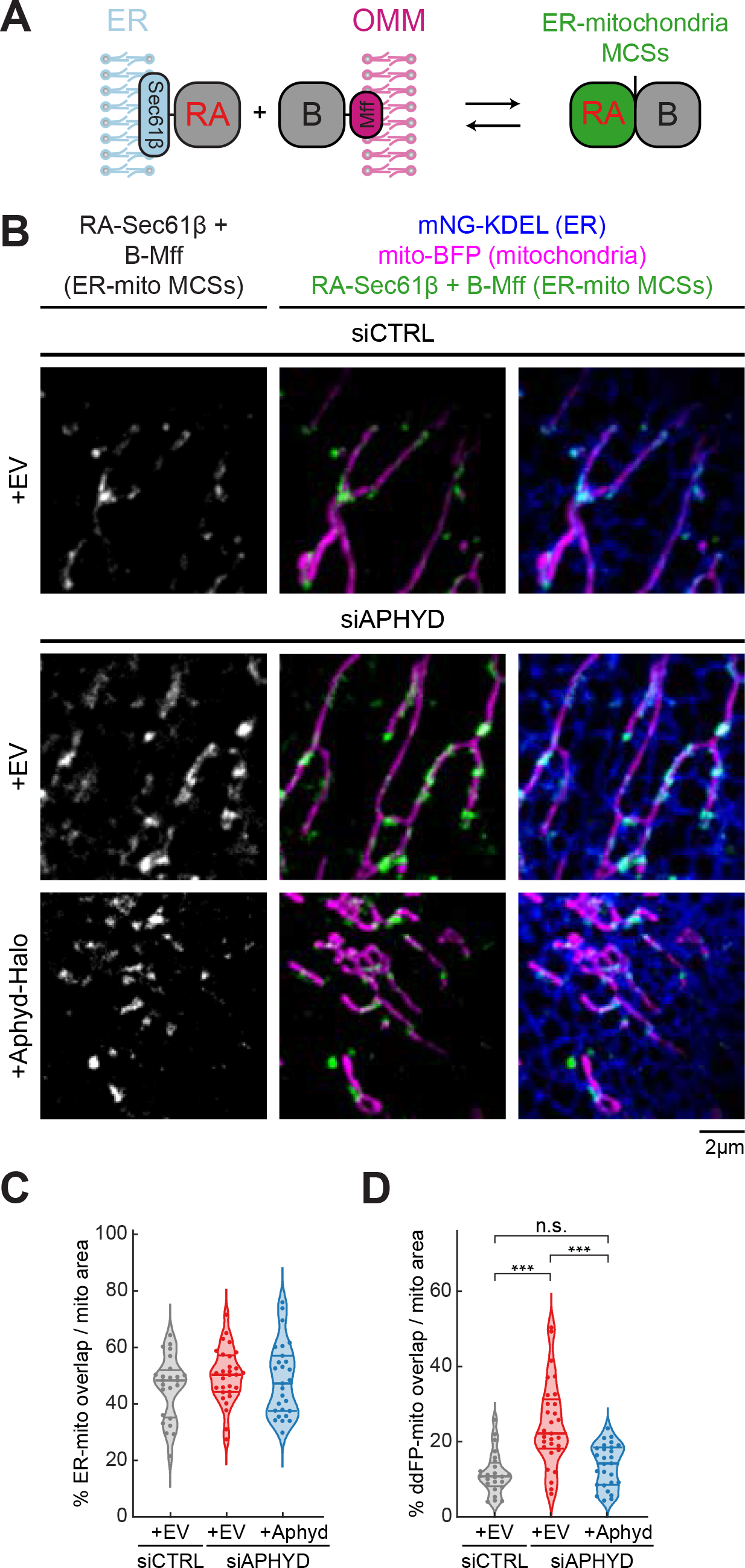
Aphyd is not required for ER-mitochondria MCS formation. (**A**) Cartoon depiction of the ER-mitochondria ddFP MCS reporter system: one monomer fused to an ER protein (RA-Sec61β) and the other monomer fused to an OMM protein (B-Mff). When ER and mitochondria come within ~10-30 nm, dimers form and fluoresce red (false colored in green) at ER-mitochondria MCSs. (**B**) Representative images of ER-mitochondria MCSs in U-2 OS cells transfected with mNG-KDEL (ER, blue), mito-BFP (magenta), RA-Sec61β and B-Mff (ER-mitochondria ddFP MCS pair, green), and either control siRNA (n=24 cells, top), Aphyd siRNA (n=29 cells, middle), or Aphyd siRNA rescued with Aphyd-Halo (n=27 cells, bottom). (**C**) Percentage of ER-mitochondria overlap over total mitochondrial area calculated by Mander’s correlation coefficient (MCC) of two binary images in 20×20 μm ROIs taken from (B). (**D**) Percentage of ddFP-mitochondria overlap over total mitochondrial area calculated by Mander’s correlation coefficient (MCC) of two binary images in 20×20 μm ROIs taken from (B). All data taken from 3 biological replicates; statistical significance calculated by one-way ANOVA. n.s., not significant; ***p≤0.001. Scale bar = 2 μm. See Figure 5-source data 1.

### Aphyd is required for ER-associated mitochondrial constriction

We have previously shown that ER-associated fission and fusion occurs at ER-associated mitochondrial constriction sites or nodes (Abrisch et al., 2020; Friedman et al., 2011). Since Aphyd depletion blocks fission and fusion machinery recruitment, we wondered whether Aphyd is required for ER-associated mitochondrial constriction. U-2 OS cells were co-transfected with mito-BFP (magenta), an ER marker (mCh-Sec61β, green), and with either Aphyd siRNA or siCTRL (Figure 6A, S6B, and S6C). ER-associated mitochondrial constrictions were defined as positions where the fluorescence intensity of the mitochondrial matrix marker (mito-BFP) dipped by >40% along several z-planes (orange arrows; ER crossings with no constriction were marked by a purple arrow, Figure S6A). Aphyd-depleted cells had 3-fold fewer ER-associated mitochondrial constrictions than siCTRL-treated cells (23% versus 63%, respectively, Figure 6A, 6C, and S6B). Re-expression of siRNA-resistant Aphyd-mNG restored levels of ER-associated mitochondrial constrictions to 75% whereas an EV control (mNG-N1) did not (Figure 6A, 6C, and S6B). These data reveal that Aphyd is required for ER-associated mitochondrial membrane constriction, which explains why its depletion disrupts fission/fusion machinery recruitment and node formation.

**Figure 6.**
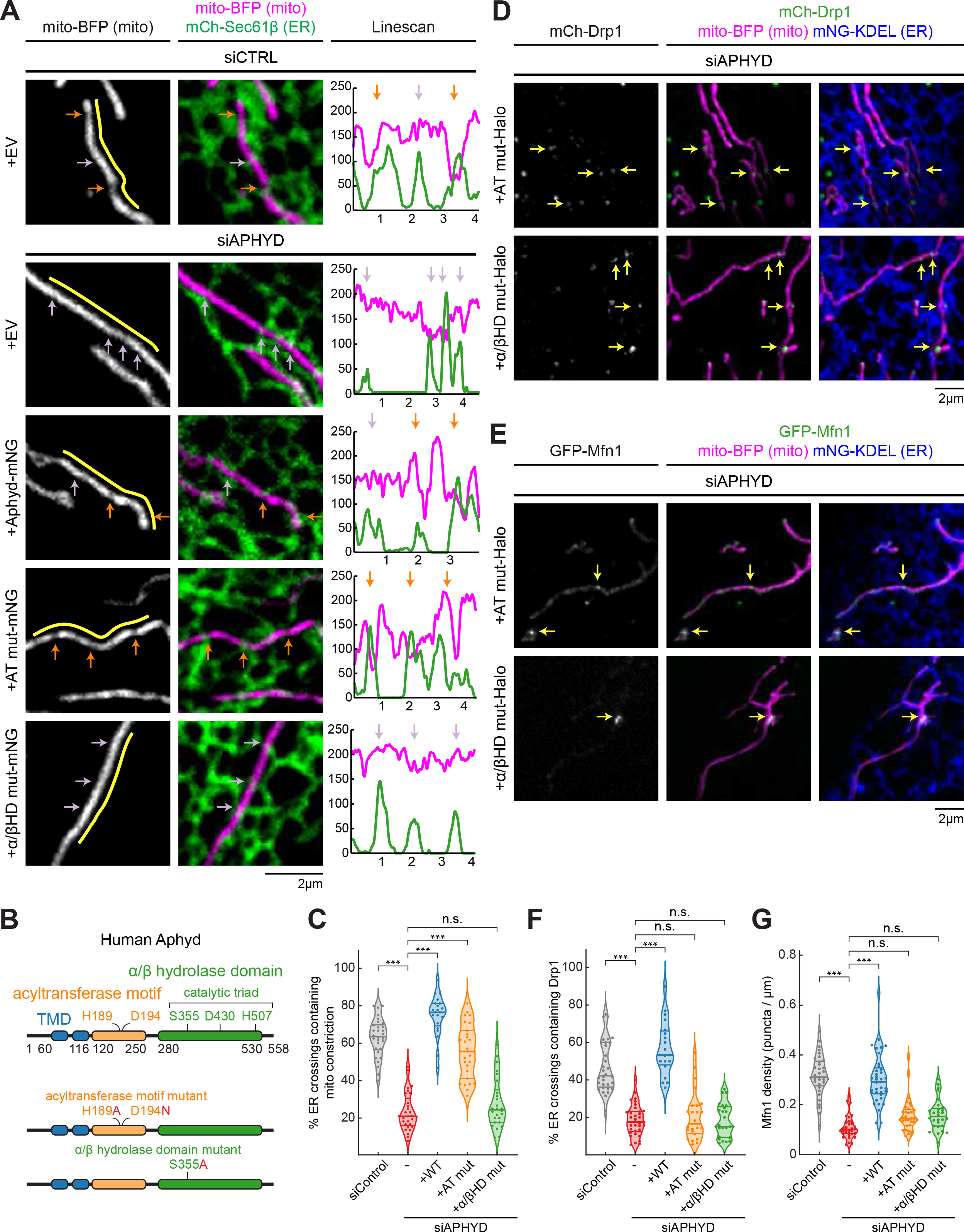
Uncoupling Aphyd’s enzymatic requirements during ER-associated mitochondrial constriction, fission, and fusion. (**A**) Representative images and line scans (yellow) of ER-mitochondria crossings in U-2 OS cells transfected with mNG-Sec61β (ER, green), mito-BFP (grey, left; magenta, middle) and either control siRNA (n=30 cells), Aphyd siRNA (n=29 cells), or Aphyd siRNA rescued with either WT Aphyd (n=27 cells), the AT mut (acyltransferase motif mutant, n=37 cells), or the α/βHD mut (alpha/beta hydrolase domain mutant, n=26 cells) mNG fusion constructs. A representative line scan (yellow, moved aside for visualization purposes) along a mitochondrion (mito-BFP) shows spatial correlation between constrictions (marked by dips in fluorescence intensity, magenta) and ER crossings (green). Orange and purple arrows mark ER crossings with and without constrictions, respectively. (**B**) Aphyd domain organization with indicated amino acids for each motif or domain (top). Mutations are indicated in red for either the AT or α/βHD mutants. (**C**) Quantification of resolvable ER crossings coincident with a mitochondrial constriction per cell from (A). (**D**) As in Figure 2B, representative images of U-2 OS cells transfected with mCh-Drp1 (green), mito-BFP (magenta), mNG-Sec61β (ER, blue), and Aphyd siRNA rescued with either the Halo-tagged Aphyd AT mut (n=24 cells, top) or α/βHD mut (n=25 cells, bottom). Yellow arrows indicate examples of Drp1 puncta at ER-mitochondria crossings. (**E**) As in Figure 2A, representative images of U-2 OS cells transfected with GFP-Mfn1 (green), mito-BFP (magenta), mCh-Sec61β (ER, blue), and Aphyd siRNA rescued with either the Halo-tagged Aphyd AT mut (n=33 cells, top) or α/βHD mut (n=28 cells, bottom). Yellow arrows indicate examples of Mfn1 puncta along mitochondria at ER-mitochondria crossings. (**F**) As in Figure 2E, quantification of percent ER crossings containing Drp1 puncta from experiments shown in (D) and Figure 2B. (**G**) As in Figure 2D, quantification of Mfn1 density along mitochondrial length from experiments shown in (E) and Figure 2A. All data taken from 3 biological replicates; statistical significance calculated by one-way ANOVA. n.s., not significant; ***p≤0.001. Scale bar = 2 μm. See Figure 6-source data 1.

### Aphyd’s alpha/beta hydrolase domain rescues mitochondrial constrictions

A local alteration in phospholipid shape at ER MCSs could change membrane curvature and might be the mechanism used by ER MCSs to constrict mitochondria (Agrawal and Ramachandran, 2019; Harayama and Riezman, 2018a; Van Meer et al., 2008). Our data have shown that Aphyd is a determinant of ER-associated mitochondrial constriction. Aphyd has two compelling and highly conserved motifs/domains predicted to alter lipid shape in opposing ways: an acyltransferase motif and an alpha/beta hydrolase domain consisting of a histidine, aspartic acid, and catalytic serine (Xu et al., 2018). Previous work suggests that the alpha/beta hydrolase domain mainly converts PS to lysoPS, while the opposing acyltransferase motif could convert single chain phospholipids, such as lysoPS, back into dual chain phospholipids (Kamat et al., 2015; Montero-Moran et al., 2010; Xu et al., 2018). We hypothesized that these enzymatic activities could alter phospholipid shape to promote ER-associated mitochondrial constriction. We generated point mutants within each of these motifs that would be predicted to abrogate enzymatic function: H189A and D194N in the acyltransferase motif (AT mut), and S355A in the alpha/beta hydrolase domain (α/βHD mut) (Figure 6B). Cells were depleted with Aphyd siRNA and co-transfected with mito-BFP (magenta) and mCh-Sec61β (ER, green) to score ER-associated mitochondrial constriction as before (Figure 6A and S6B). SiRNA-resistant WT and mutant Aphyd constructs were tested for their ability to rescue ER-associated mitochondrial constriction. The acyltransferase motif mutant could rescue ER-associated mitochondrial constriction to a level similar to WT rescue levels, whereas the alpha/beta hydrolase domain mutant could not (Figure 6A, 6C, S6B, and S6C). These data show that phospholipid hydrolysis from a dual chain phospholipid to a single chain phospholipid is part of the mechanism by which Aphyd drives ER-associated mitochondrial constriction.

### Aphyd’s acyltransferase and hydrolase domain are required for fission and fusion

Since the alpha/beta hydrolase activity is sufficient to rescue ER-associated mitochondrial constriction (measured with mito-BFP fluorescence intensity), we expected that it might also rescue fission and fusion node formation. First, we scored Aphyd motif/domain requirements for fission (Drp1) machinery recruitment to ER-mitochondria MCSs. Each mutant was tested for its ability to rescue Drp1 recruitment in cells depleted of endogenous Aphyd with siRNA. These cells were co-transfected with siRNA-resistant mutant Aphyd along with mito-BFP (magenta), mCh-Sec61β (ER, blue), and mCh-Drp1 (green) (Figure 6D, S6D, and S6H). Interestingly, only WT Aphyd could restore Drp1 recruitment to ER crossings (Figure 2B, 2E, 6D, 6F, and S6D). Neither the acyltransferase motif nor alpha/beta hydrolase domain mutants could rescue Drp1 recruitment in Aphyd-depleted cells (Figure 6D, 6F, and S6D). In complementary experiments, we also tested whether each mutant could rescue Mfn1 fusion machinery recruitment. Cells were co-transfected with either a control or Aphyd siRNA, fluorescently tagged Mfn1 (GFP-Mfn1, green), a mito-BFP (magenta), and mNG-Sec61β (ER, blue) and were rescued with siRNA-resistant WT or mutant Aphyd constructs (Aphyd-Halo) (Figure 6E and S6E). Similar to what was seen for Drp1, only the WT protein could restore Mfn1 accumulation at ER-mitochondria MCSs in Aphyd-depleted cells (Figure 2A, 2D, 6E, and 6G). Thus, both acyltransferase and alpha/beta hydrolase activities are required for the recruitment of fission and fusion machinery to ER-associated nodes. Consistent with these results, only the WT Aphyd can rescue overall mitochondrial morphology (Figure S6F and S6G). So, while the alpha/beta hydrolase domain is sufficient for ER-associated mitochondrial constriction, the acyltransferase motif is still required to form active nodes for fission and fusion. These data suggest that the mechanism of ER-associated mitochondrial node formation is a multistep process that requires reversible modifications of phospholipids coming from the ER.

### ORP8 is required to deliver Aphyd-induced altered phospholipids to mitochondria

Since ER-localized Aphyd is necessary to sustain mitochondrial fission and fusion, we identified potential candidates that could facilitate lipid exchange during Aphyd-driven mitochondrial node formation. Two published ER-localized lipid transfer proteins were worthy candidates: ORP8, which has been proposed to transfer phospholipids, such as PS, to other organelles at ER-mitochondria and ER-PM MCSs (Chung et al., 2015; Galmes et al., 2016) and the VPS13A/VPS13D paralogs, which have demonstrated roles in facilitating lipid transfer between the ER and other organelles at MCSs (Guillén-Samander et al., 2021a; Kumar et al., 2018). To score whether these ER-localized lipid transport proteins contribute to steady state mitochondrial morphology, U-2 OS cells were co-transfected with a mitochondrial matrix marker (mito-BFP, magenta) and either control (siCTRL), OSBPL8 siRNA, or VPS13A/VPS13D siRNAs (which led to efficient protein depletion, Figure 7F and 7G) and mitochondrial morphology was quantified as described previously (Lee et al., 2016). Neither ORP8 depletion nor VPS13A/13D depletion by siRNA treatment altered mitochondrial morphology compared to siCTRL-treated cells (Figure 7A, 7C, 7D, and 7E). We next tested whether ORP8 or VPS13A/D were required for the dramatic mitochondrial fragmentation phenotype observed upon Aphyd OE. Cells were transfected with either siCTRL, OSBPL8 siRNA, or VPS13A/VPS13D siRNAs and with either EV control (mCherry-N1) or Aphyd-mCh, as indicated (Figure 7A, 7B, and 7C). Aphyd-mCh OE promotes mitochondrial fragmentation as previously described (Figure 4F, 7B, and 7C). However, mitochondrial morphology is dramatically rescued upon ORP8 depletion and re-fragmented by exogenous siRNA-resistant ORP8 re-expression in the AphydOE/ORP8-depleted cells (Figure 7B and 7D). In contrast, depletion of both VPS13A and VPS13D did not prevent Aphyd OE-induced fragmentation (Figure 7C and 7E). Thus, ORP8, but not VPS13A/D, facilitates Aphyd activity during ER-associated mitochondrial constriction. A likely hypothesis is that Aphyd and ORP8 function to alter the lipid composition at ER-associated mitochondrial constrictions for fission and fusion machinery recruitment.

**Figure 7.**
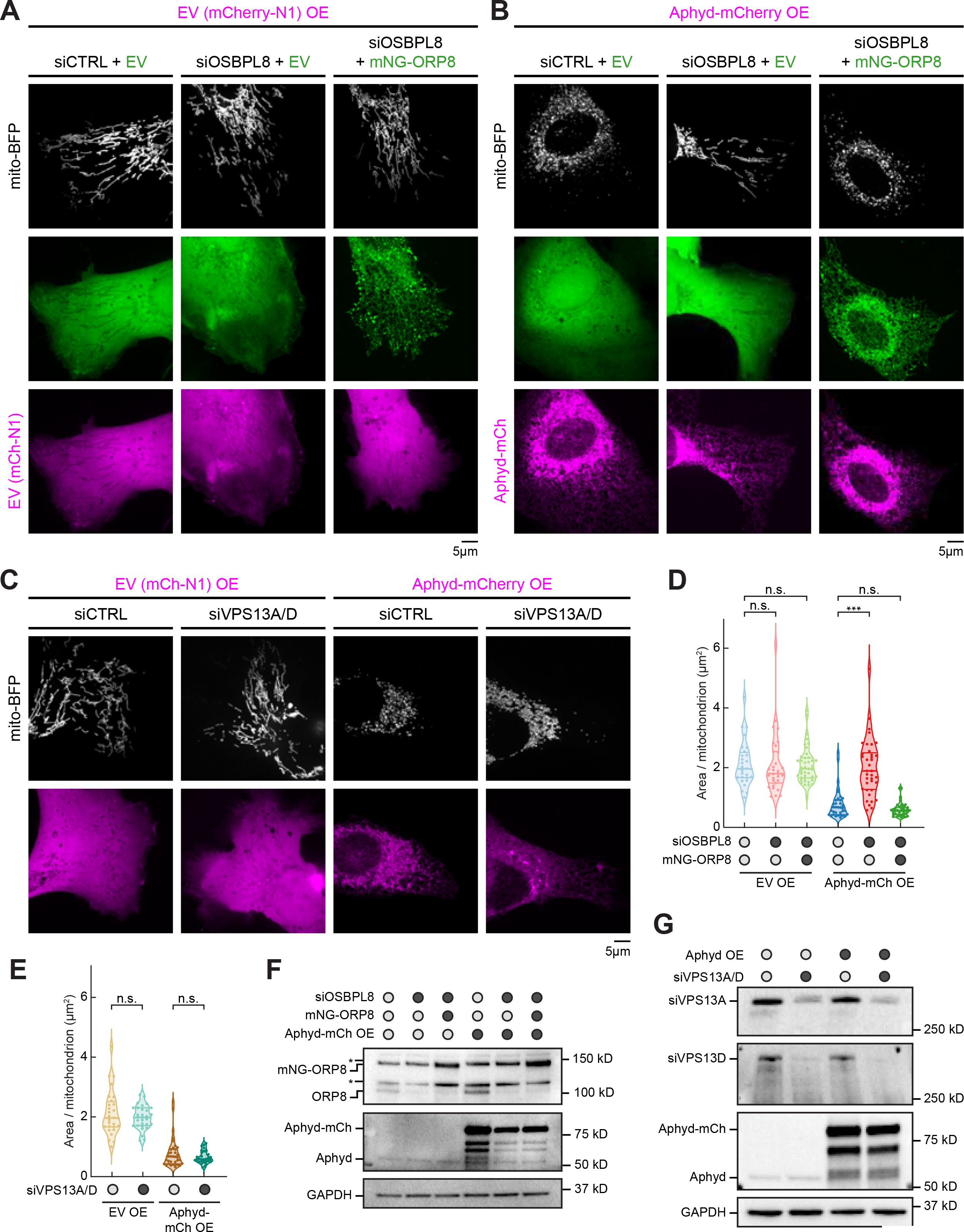
ORP8 is required to deliver Aphyd-induced altered phospholipids to mitochondria. (**A**) Representative images of mitochondrial morphology (labeled by mito-BFP, grey) of U-2 OS cells transfected with EV (mCherry-N1, magenta) and with either siCTRL and mNG-C1 EV (green, n=28 cells, left); OSBPL8 siRNA and mNG-C1 EV (green, n=21 cells, middle); or OSBPL8 siRNA and siRNA-resistant mNG-ORP8 (green, n=41 cells, right). (**B**) Representative images of mitochondrial morphology (labeled by mito-BFP, grey) of U-2 OS cells transfected with Aphyd-mCherry OE (magenta) and either siCTRL and mNG-C1 EV (green, n=35 cells, left); OSBPL8 siRNA with mNG-C1 EV (green, n=37 cells, middle); or OSBPL8 siRNA with siRNA-resistant mNG-ORP8 (green, n=37 cells, right). (**C**) Representative images of mitochondrial morphology (labeled by mito-BFP, grey) of U-2 OS cells transfected with either EV (mCherry-N1, magenta) and siCTRL (n=28 cells); EV and VPS13A/D siRNA (n=30 cells); with Aphyd-mCherry OE (magenta) and siCTRL (n=36 cells); or with Aphyd-mCherry OE (magenta) and VPS13A/D siRNA (n=29 cells). (**D**) Quantification of mean mitochondrial size (area per mitochondrion in μm^2^) within a 15×15 μm ROI from (A) and (B). (**E**) Quantification of mean mitochondrial size (area per mitochondrion in μm^2^) within a 15×15 μm ROI from (C). (**F**) Representative immunoblot shows efficiency of depletion in U-2 OS cells from (A) and (B) treated with control siRNA or OSBPL8 siRNA and rescued with siRNA-resistant mNG-ORP8. GAPDH serves as a loading control. Asterisk indicates non-specific band. (**G**) Representative immuno-blot shows efficiency of depletion in U-2 OS cells from (C) treated with control siRNA or VPS13A/D siRNA. GAPDH serves as a loading control. All data taken from 3 biological replicates; statistical significance calculated by one-way ANOVA. n.s., not significant; ***p≤0.001. Scale bar = 5 μm. See Figure 7-source data 1-9.

## Discussion

Through proximity proteomics, we have identified an ER membrane protein, Aphyd that regulates the formation of fission and fusion nodes at ER-mitochondria MCSs. Aphyd localizes to the ER membrane and to a lesser extent mitochondria. Interestingly, only the ER-localized form is necessary and sufficient to maintain mitochondrial morphology. We have shown that Aphyd is required for mitochondrial constrictions prior to the formation of fission and fusion nodes. Without Aphyd, nodes are not restored, leading to an overall elongated morphology due to overruling tip-to-tip fusion. Two critical motifs/domains required for these activities are the alpha/beta hydrolase domain, which converts dual chain phospholipids to lysophospholipids, and the acyltransferase motif, which converts the inverse reaction. We have identified that only the alpha/beta hydrolase domain is required for inner mitochondrial membrane constriction. However, the acyltransferase motif is required for a step downstream of IMM constrictions to allow fission and fusion machineries to be recruited to the OMM. Indeed, both the alpha/beta hydrolase domain and acyltransferase motif are required to restore mitochondrial morphology.

We propose a model whereby Aphyd’s alpha/beta hydrolase domain first promotes single chain phospholipid build up along the mitochondrial membrane and then the acyltransferase motif promotes dual chain phospholipid build up. Both single chain and dual chain phospholipids are required for positive and negative membrane curvature respectively, which are required for mitochondrial efficient fission and fusion processes (Figure 8, Model). We have shown that ER-localized Aphyd is necessary and sufficient to restore ER-associated mitochondrial morphology and that Aphyd functions on the ER to affect mitochondrial fission and fusion cycles.

**Figure 8.**
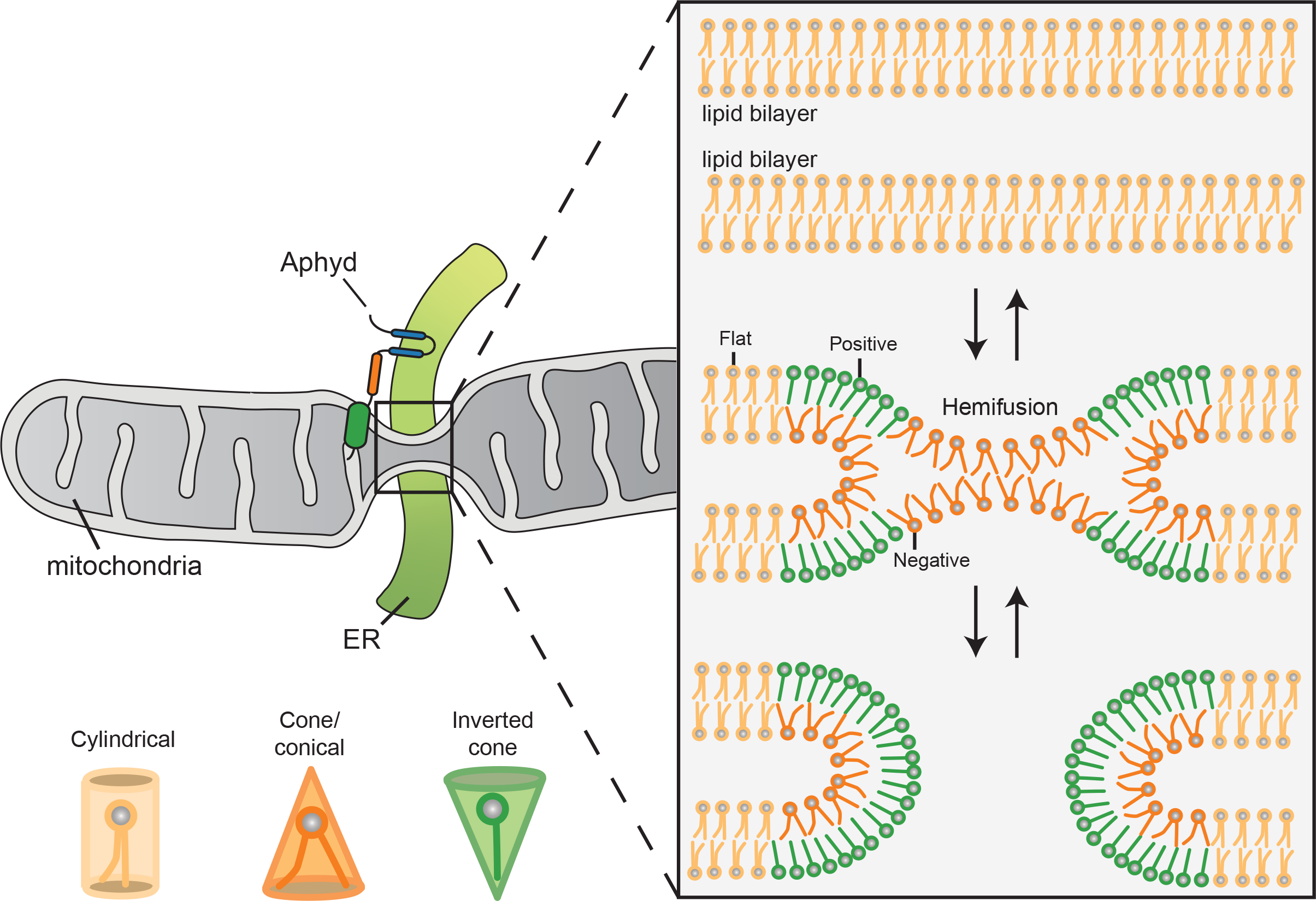
Model for how Aphyd promotes both lysophospholipid and phospholipid formation required for positive and negative membrane curvature to support efficient cycles of fission and fusion.

We have always considered that lipid composition and shape is likely a contributor to protein localization and membrane architecture at MCSs but by what mechanism was unclear. Lipid composition can modulate physicochemical properties of organelle membranes and shape due to the diversity of each phospholipid head group and hydrophobic tail (Harayama and Riezman, 2018b). For example, phosphatidylethanolamine (PE) exists as a cone-shaped lipid, which can create negative spontaneous curvature to promote hemi-fusion intermediates. Cone-shaped lipids have been showed to be required for SNARE-mediated fusion and osteoclast fusion (Irie et al., 2017; Zick et al., 2014). These data fall in line with Aphyd converting phospholipids to lysophospholipids and vice versa to create spontaneous negative and positive membrane curvature (respectively) necessary for hemi-fusion intermediates at MCSs for constriction, fission, and fusion.

Additionally, many known mitochondrial fission and fusion factors have been associated with specific phospholipids. *In vitro* and cryoEM studies of Drp1 showed that reconstituted Drp1 prefers to oligomerize around negatively charged phospholipids such as PS and cardiolipin (CL) (Francy et al., 2017; Kalia et al., 2018). Additionally, *in vitro* reconstitution of OPA1 and CL membranes are sufficient for tethering and fusion (Ban et al., 2017). Interestingly, Aphyd has the highest specificity for phosphotidylserine (PS) (Kamat et al., 2015), which constitutes only about 1-2% of the total mitochondrial membrane phospholipid composition (Van Meer et al., 2008). Thus, it is likely that PS and lysoPS conversion could be secluded to the ER-mitochondrial membrane contacts sites deemed competent for constriction and fission/fusion machinery recruitment. Intriguingly, it is unclear how the adaptor proteins, Mff, Mid49, and Mid51 enrich in punctate localizations at ER-mitochondria contacts prior to Drp1 localization (Friedman et al., 2011; Palmer et al., 2011). Perhaps lipid alteration and the curvature sensing portions of these adaptor proteins could be responsible for its punctate localization and it is Aphyd that creates these lipid changes.

In addition, the ER and mitochondria also form contact sites to transfer phospholipids (Achleitner et al., 1999; Ardail et al., 1993; Kornmann et al., 2009; Vance, 1990). Several studies have shown that lipid transfer and mitochondrial morphology are highly interdependent. More specifically, in yeast, the ER-mitochondria encounter structure (ERMES) complex, which is proposed to transfer ER PS and PC to mitochondrial membranes, (Jeong et al., 2017; Kojima et al., 2016; Kornmann et al., 2009) assembles between the ER and mitochondrial membranes at MCSs (Michel and Kornmann, 2012; Murley et al., 2013). Interestingly, a deletion in the single ERMES complex protein, Gem1, also fragments mitochondria (Frederick et al., 2004; Murley et al., 2013). In animal cells, Gem1 homologs are Miro1 and Miro2. Miro1/2 proteins regulate mitochondrial and peroxisomal trafficking on microtubules (MTs) and can elongate both mitochondria and peroxisomes (Castro et al., 2018; Fransson et al., 2006; MacAskill et al., 2009b, 2009a; Modi et al., 2019; Okumoto et al., 2018; Russo et al., 2009; Saotome et al., 2008). The De Camilli lab has shown that the lipid transport protein VPS13D is recruited by Miro (Guillén-Samander et al., 2021b) suggesting that lipid transfer must be required for lipid membrane expansion and subsequent elongated mitochondrial morphology. Additionally, recent work from the Nunnari lab also suggests that a buildup of lysophosphatidic acid (LPA), which promotes positive membrane curvature, sensed by mitochondrial carrier homologue 2 (MTCH2) stimulates mitochondrial fusion (Labbé et al., 2021). These data are intriguing because Aphyd could also promote positive membrane curvature for the formation of fission and fusion nodes. Taken together, these data suggest that lipid transfer and modification at ER-mitochondria MCSs has a direct involvement with mitochondrial elongation and fragmentation by affecting fusion and fission.

## Acknowledgements

We thank Jonathan Friedman, Rob Abrisch, Robert Campbell, David Sabatini, and Alice Ting for sharing plasmids. We thank Luke Lavis for sharing Janelia Fluor 646 SE and Janelia Fluor X 646 for SNAP and Halo tag imaging. We thank Alex Rosa Campos for collecting and analyzing mass spectrometry data. We thank Jonathan Friedman, Eric Sawyer, Jonathan Striepen, and Sofia Zaganelli for experimental suggestions. T.T.N. was supported by grants from the NIH (T32 GM008759 and T32 GM142607; GM120998 to G.K.V.). G.K.V. is an investigator of the Howard Hughes Medical Institute.

## Author contributions

T.T.N. and G.K.V. contributed to experimental design. T.T.N. conducted all experiments, data analysis, and figure composition. T.T.N. wrote the manuscript. G.K.V. edited the manuscript.

## Data and material availability

All data is available in the main text or the supplementary materials.

## Supplemental information

**Figure S1.**
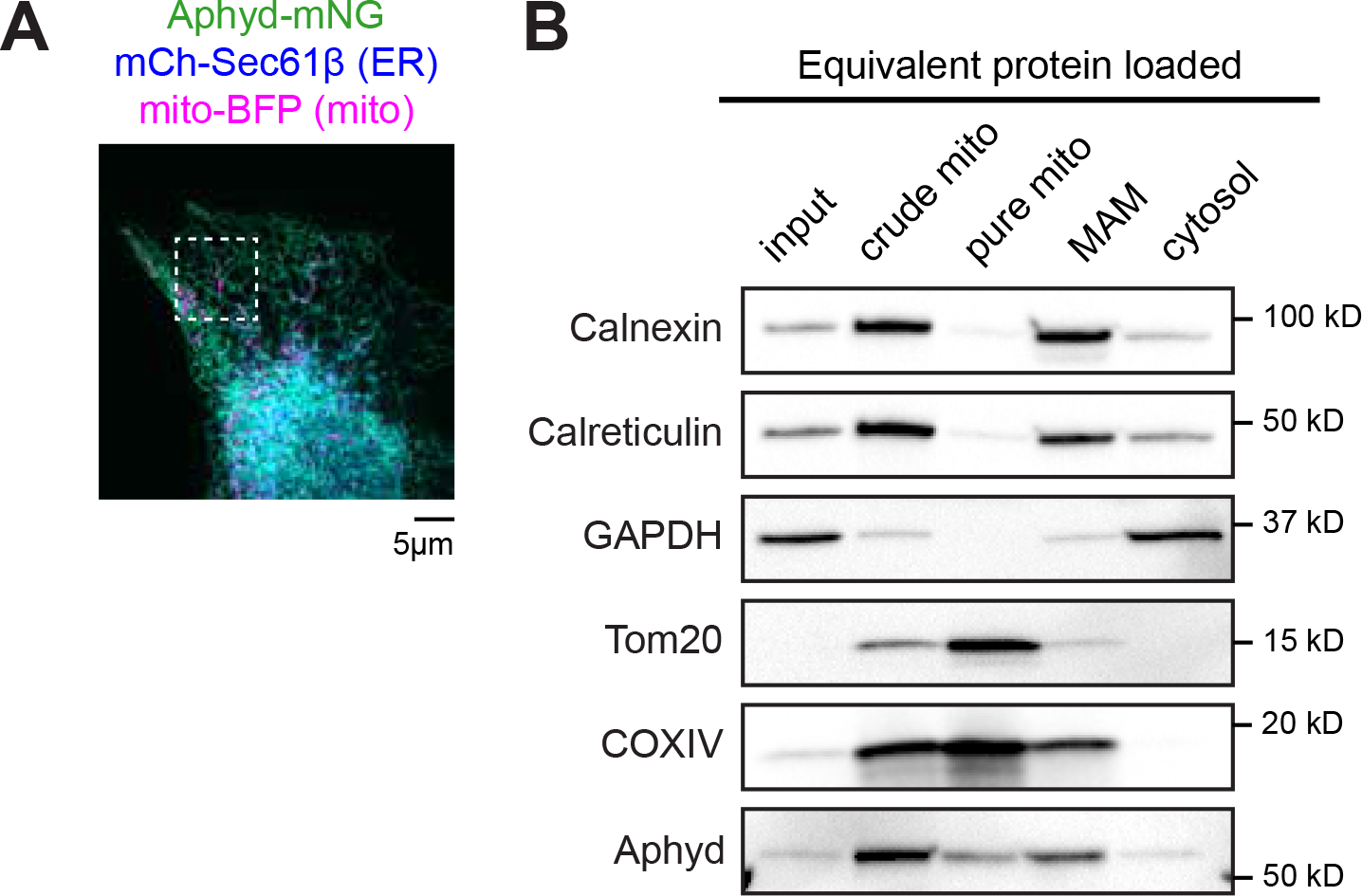
Endogenous Aphyd localizes to the ER and mitochondria, to a lesser extent (related to Figure 1). (**A**) Representative image from Figure 1I of a U-2 OS cell expressing low levels of Aphyd-mNG (green), mCh-Sec61β (ER, blue), and mito-BFP (mitochondria, magenta). White dashed box indicates inset in Figure 1I. (**B**) Immunoblot of various membrane markers after dounce homogenization, differential centrifugation, and Ficoll mitochondrial purification, yielding a crude mitochondrial fraction (prior to Ficoll spin), pure mitochondrial fraction (after Ficoll spin), mitochondrial associated membrane (MAM) fraction (after Ficoll spin), and cytosol (containing ER and other light membrane fractions). ER = Calnexin, calreticulin. Cytosol = GAPDH. Tom20, COXIV = mitochondria. Scale bar = 5 μm. See Figure 1-figure supplement 1-source data 1-6.

**Figure S2.**
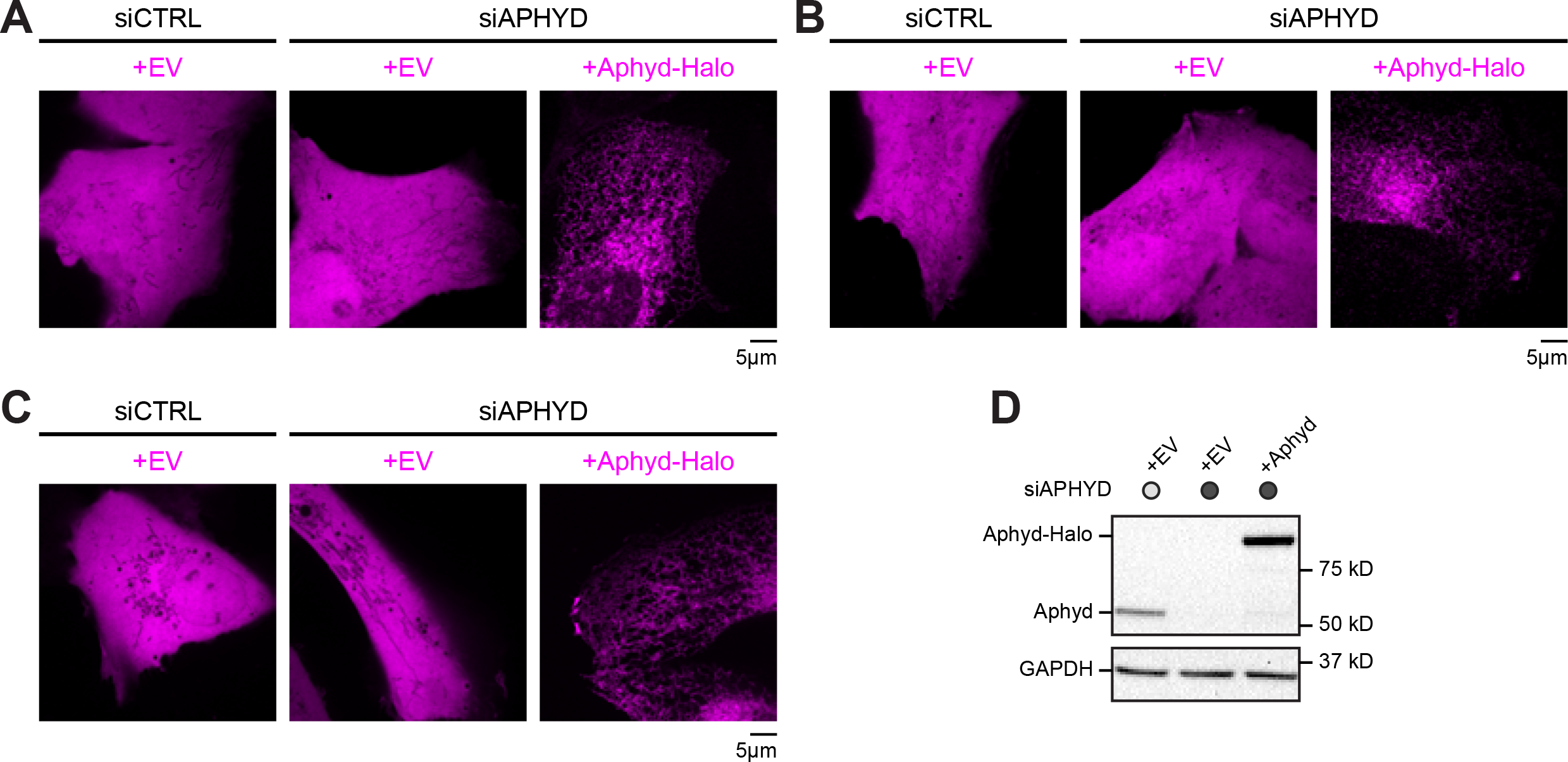
Aphyd depletion decreases fusion/fission node formation (related to Figure 2). (**A**) Representative images of either EV (Halo-N1) or Halo-tagged expression constructs (magenta) visualized using JF X 646 corresponding to cells from Figure 2A. (**B**) Representative images of either EV (Halo-N1) or Halo-tagged expression constructs (magenta) visualized using JF X 646 corresponding to cells from Figure 2B. (**C**) Representative images of either EV (Halo-N1) or Halo-tagged expression constructs (magenta) visualized using JF X 646 corresponding to cells from Figure 2C. (**D**) Representative immunoblot shows efficiency of depletion in U-2 OS cells from Figure 2 treated with control siRNA or Aphyd siRNA and rescued with siRNA-resistant Aphyd-Halo. GAPDH serves as a loading control. All data taken from 3 biological replicates. Scale bar = 5 μm. See Figure 2-figure supplement 1-source data 1-3.

**Figure S3.**
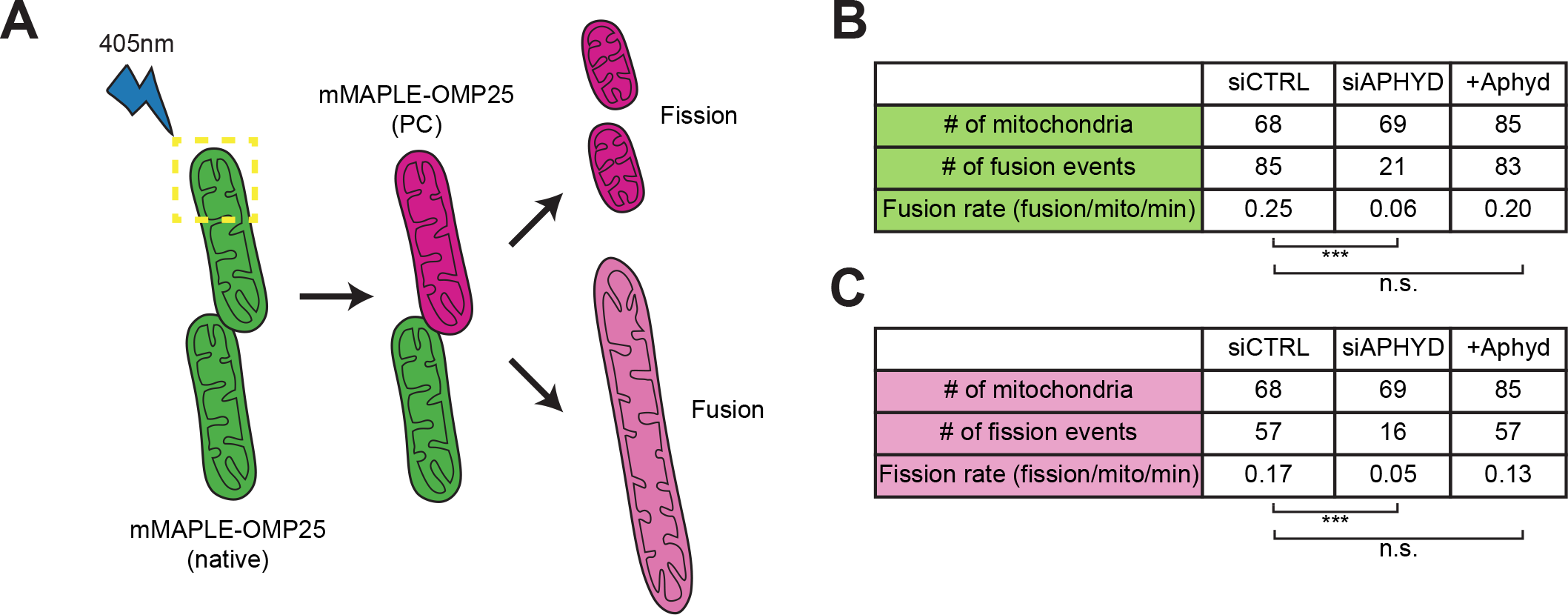
Aphyd depletion reduces fusion and fission rate (related to Figure 3). (**A**) Cartoon demonstrating mitochondria labeled with mMAPLE-OMP25 (green) undergoing photoconversion upon 405 nm light exposure, converting mMAPLE to red (magenta). Upon fission or fusion, red fluorescence separates or diffuses. (**B**) Expanded table from Figure 3D for fusion rate per mitochondrion per minute from experiments shown in Figure 3A-C. (**C**) Expanded table from Figure 3E for fission rate per mitochondrion per minute from experiments shown in Figure 3A-C. All data taken from 3 biological replicates; statistical significance calculated by one-way ANOVA. n.s., not significant; ***p≤0.001.

**Figure S4.**
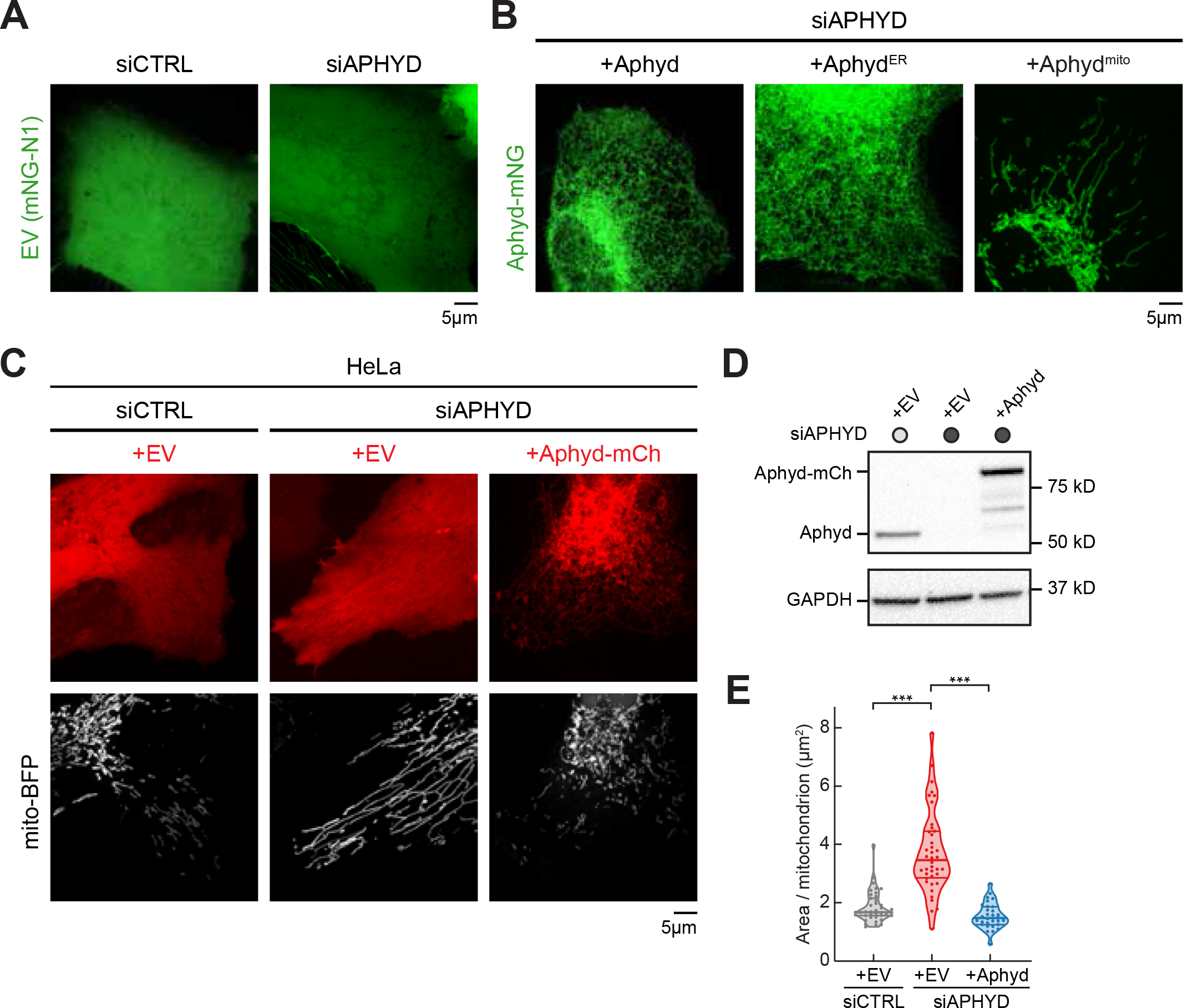
Aphyd is required to maintain mitochondrial morphology in HeLa cells (related to Figure 4). (**A**) Representative images of EV constructs (green) corresponding to cells from Figure 4B. (**B**) Representative images of mNG-tagged expression constructs (green) corresponding to cells from Figure 4C. (**C**) Representative images of mitochondrial morphology (labeled by mito-BFP, grey) of HeLa cells transfected with control siRNA (n=39 cells, left), Aphyd siRNA (n=40 cells, middle), or Aphyd siRNA rescued with siRNA-resistant Aphyd-mCherry construct (n=30 cells, right). (**D**) Immunoblot shows efficiency of depletion in HeLa cells from (C) treated with control siRNA or Aphyd siRNA and rescued with siRNA-resistant Aphyd-mCherry. GAPDH serves as a loading control. (**E**) Quantification of mean mitochondrial size (area per mitochondrion in μm^2^) within a 15×15 μm ROI from (C). All data taken from 3 biological replicates; statistical significance calculated by one-way ANOVA. ***p≤0.001. Scale bar = 5 μm. See Figure 4-figure supplement 1-source data 1-4.

**Figure S5.**
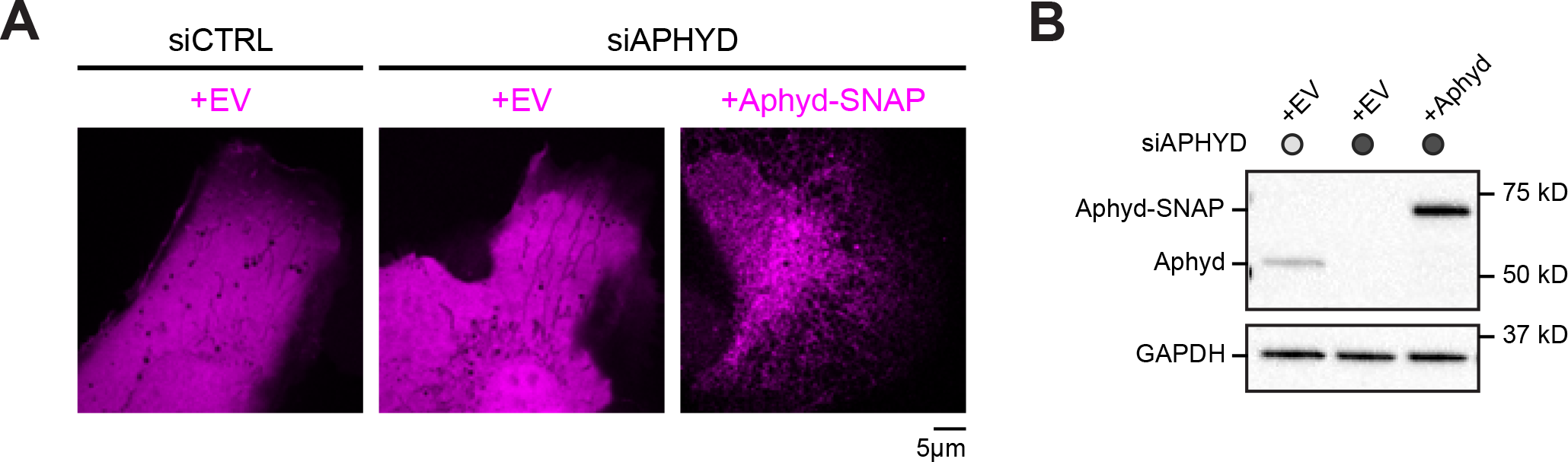
Aphyd is not required for ER-mitochondria MCS formation (related to Figure 5). (**A**) Representative images of either EV (SNAP-N1) or SNAP-tagged expression constructs (magenta) visualized using JF 646, SE corresponding to cells from Figure 5B. (**B**) Representative immunoblot shows efficiency of depletion in U-2 OS cells from Figure 5B treated with control siRNA or Aphyd siRNA and rescued with siRNA-resistant Aphyd-SNAP. GAPDH serves as a loading control. Scale bar = 5 μm. See Figure 5-figure supplement 1-source data 1-3.

**Figure S6.**
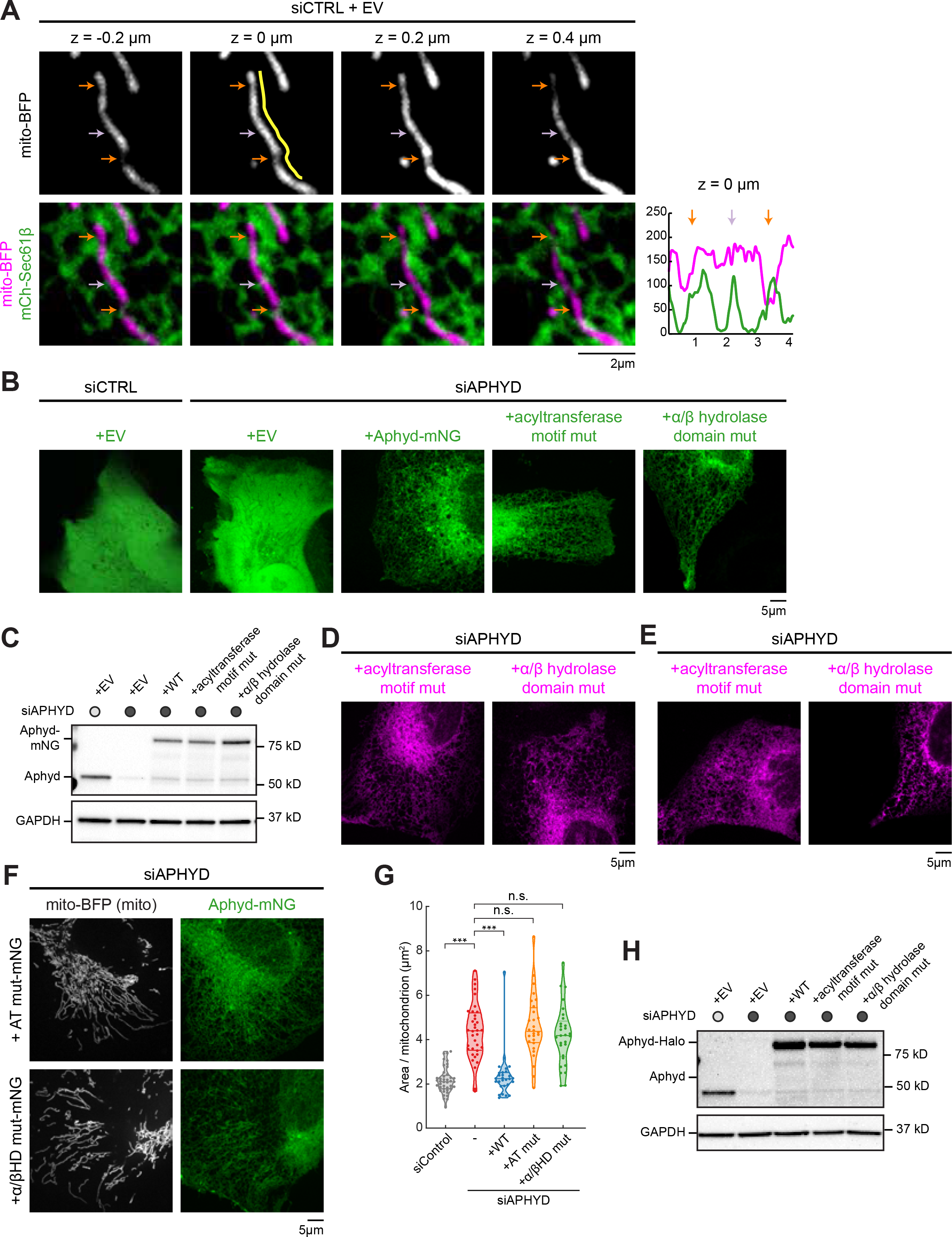
Aphyd’s acyltransferase motif and alpha/beta hydrolase domain are required for mitochondrial morphology (related to Figure 6). (**A**) Representative z-series and line scans (yellow) of ER-mitochondria crossings in U-2 OS cell from Figure 6A transfected with mCh-Sec61β (ER, green), mito-BFP (grey, top; magenta, bottom), and an EV (not shown). Images display examples of mitochondrial focal planes chosen for line scan analysis (z = 0 μm). Constrictions at ER crossings are indicated by orange arrows. ER-mitochondria crossings containing no constrictions are indicated by purple arrows. (**B**) Representative images of either EV (mNG-N1) or mNG-tagged expression constructs (green) corresponding to cells from Figure 6A. (**C**) Immunoblot shows efficiency of depletion in U-2 OS cells from Figure 6A treated with control siRNA or Aphyd siRNA and rescued with siRNA-resistant Aphyd-mNG constructs. GAPDH serves as a loading control. (**D**) Representative images of Halo-tagged expression constructs (magenta) visualized using JF X 646 corresponding to cells from Figure 6D. (**E**) Representative images of Halo-tagged expression constructs (magenta) visualized using JF X 646 corresponding to cells from Figure 6E. (**F**) Representative images of mitochondrial morphology (labeled by mito-BFP, grey) of U-2 OS cells transfected with Aphyd siRNA and either the acyltransferase motif mutant (AT mut, n=29 cells, top) or the alpha/beta hydrolase domain mutant (α/βHD mut, n=28 cells, bottom). (**G**) Quantification of mean mitochondrial size (area per mitochondrion in μm^2^) within a 15×15 μm ROI from (F) and Figure 4D. (**H**) Representative immunoblot shows efficiency of depletion in U-2 OS cells from Figure 6D and 6E treated with control siRNA or Aphyd siRNA and rescued with siRNA-resistant Aphyd-Halo constructs. GAPDH serves as a loading control. All data taken from 3 biological replicates; statistical significance calculated by one-way ANOVA. n.s., not significant; ***p≤0.001. Scale bar = 2 μm and 5 μm, respectively. Figure 6-figure supplement 1-source data 1-7.

## Materials and methods

### DNA plasmids and primer sequences

V5-TurboID-NES was a gift from Alice Ting (Addgene plasmid # 107169) (Branon et al., 2018). V5-TurboID-Mfn1 and V5-TurboID-Mfn1E209A were generated by subcloning TurboID from V5-TurboID-NES and AgeI/XhoI sites were used to replace GFP in GFP-Mfn1 or GFP-Mfn1E209A (Abrisch et al., 2020). GFP-Mfn1 and GFP-Mfn1E209A were previously described (Abrisch et al., 2020). Mito-BFP was previously described (Friedman et al., 2011). Mitochondrial targeting sequence of budding yeast COX4 gene (aa 1-22) were PCR amplified and cloned with XhoI/BamHI sites into mTagBFP-N1. ABHD16A, renamed to Aphyd, (isoform a, NM_021160.3) was PCR amplified from HeLa cDNA and cloned into HindIII/SacII sites of mNeonGreen-N1 (purchased from Allele Biotechnology) to generate Aphyd-mNG. mCherry-Sec61β (Zurek et al., 2011) subcloned from AcGFP-Sec61β (Shibata et al., 2008) into mCherry-C1 using BglII/EcoRI sites. Stim1NTD-Aphydcyto-mNG (Aphyd^ER^) and Tom70NTD-Aphydcyto-mNG (Aphyd^mito^) were generated with Gibson assembly following manufacturer’s protocol (NEB). The templates were Aphyd-mNG and HeLa cDNA for Stim1’s NTD and Tom70’s NTD. Primers used to generate these chimeric constructs are: TN378-TN381 for Stim1NTD-Aphydcyto-mNG and Tom70NTD-Aphydcyto-mNG. SiRNA-resistant Aphyd constructs were generated from Aphyd-mNG by QuikChange II site-directed mutagenesis (Aligent Technologies Cat. #200524) following manufacturer’s protocol. Primers used were: TN361 and TN362. SiRNA-resistant Aphyd-mCherry was subcloned using HindIII/SacII sites into mCherry-N1 from siRNA-resistant Aphyd-mNG. mCherry-Drp1 was previously described (Friedman et al., 2011). Drp1 (NM_005690) was PCR amplified from HeLa cDNA and cloned into XhoI/BamHI sites substituting α-Tubulin in mCherry-α-Tubulin. mNG-Sec61β was generated by PCR amplifying human Sec61β from AcGFP-Sec61β (Shibata et al., 2008) and inserted into mNG-C1 (purchased from Allele Biotechnology) using HindIII/KpnI sites. mNG-Mfn1 was subcloned using EcoRI/BamHI sites into mNG-N1 from GFP-Mfn1. SiRNA-resistant Aphyd-Halo was subcloned using HindIII/SacII sites into Halo-N1 from siRNA-resistant Aphyd-mNG. mMAPLE-OMP25 was a gift from R. Abrisch. mMAPLE was PCR amplified from mito-mMAPLE and mMAPLE-C1 vector was derived from AcGFP-C1 by replacing GFP using NheI/BspEI sites. OMP25 was PCR amplified from paGFP-Omp25 (gift from D. Sabatini, Addgene plasmid #69598) and cloned with XhoI/BamHI into mMaple-C1. RA-Sec61β was previously described (Abrisch et al., 2020). RA was PCR amplified from RA-NES (gift from R. Campbell, Addgene plasmid #61019) (Ding et al., 2015) and RA-C1 vector was derived from AcGFP-C1 by replacing GFP using NheI/BspEI sites. Sec61β was PCR amplified and cloned into XhoI/KpnI sites of RA-C1. B-Mff was previously described (Abrisch et al., 2020). GB was PCR amplified from GB-NES (gift from R. Campbell, Addgene plasmid #61017) (Ding et al., 2015) and cloned into NheI/BspEI sites by replacing GFP in GFP-Mff [Addgene plasmid #49153, (Friedman et al., 2011)]. SiRNA-resistant Aphyd-SNAP was subcloned using HindIII/SacII sites into SNAP-N1 from siRNA-resistant Aphyd-mNG. SiRNA-resistant Aphyd-mNG acyltransferase motif mutant (H189AD194N) and alpha/beta hydrolase domain mutant (S355A) were generated from siRNA-resistant Aphyd-mNG by QuikChange II site-directed mutagenesis (Aligent Technologies Cat. #200524) following manufacturer’s protocol. Primers used were: TN329-TN332 for each mutation. SiRNA-resistant Aphyd-Halo acyltransferase motif mutant (H189AD194N) and alpha/beta hydrolase domain mutant (S355A) was subcloned using HindIII/SacII sites into Halo-N1 from siRNA-resistant Aphyd-mNG mutants. ORP8 (isoform a, NM_020841.5) was PCR amplified from HeLa cDNA and cloned into SalI/SacII sites of mNeonGreen-C1 (purchased from Allele Biotechnology) to generate mNG-ORP8. SiRNA-resistant mNG-ORP8 was generated from mNG-ORP8 by QuikChange II site-directed mutagenesis (Aligent Technologies Cat. #200524) following manufacturer’s protocol. Primers used were: TN541-542.

### Primers

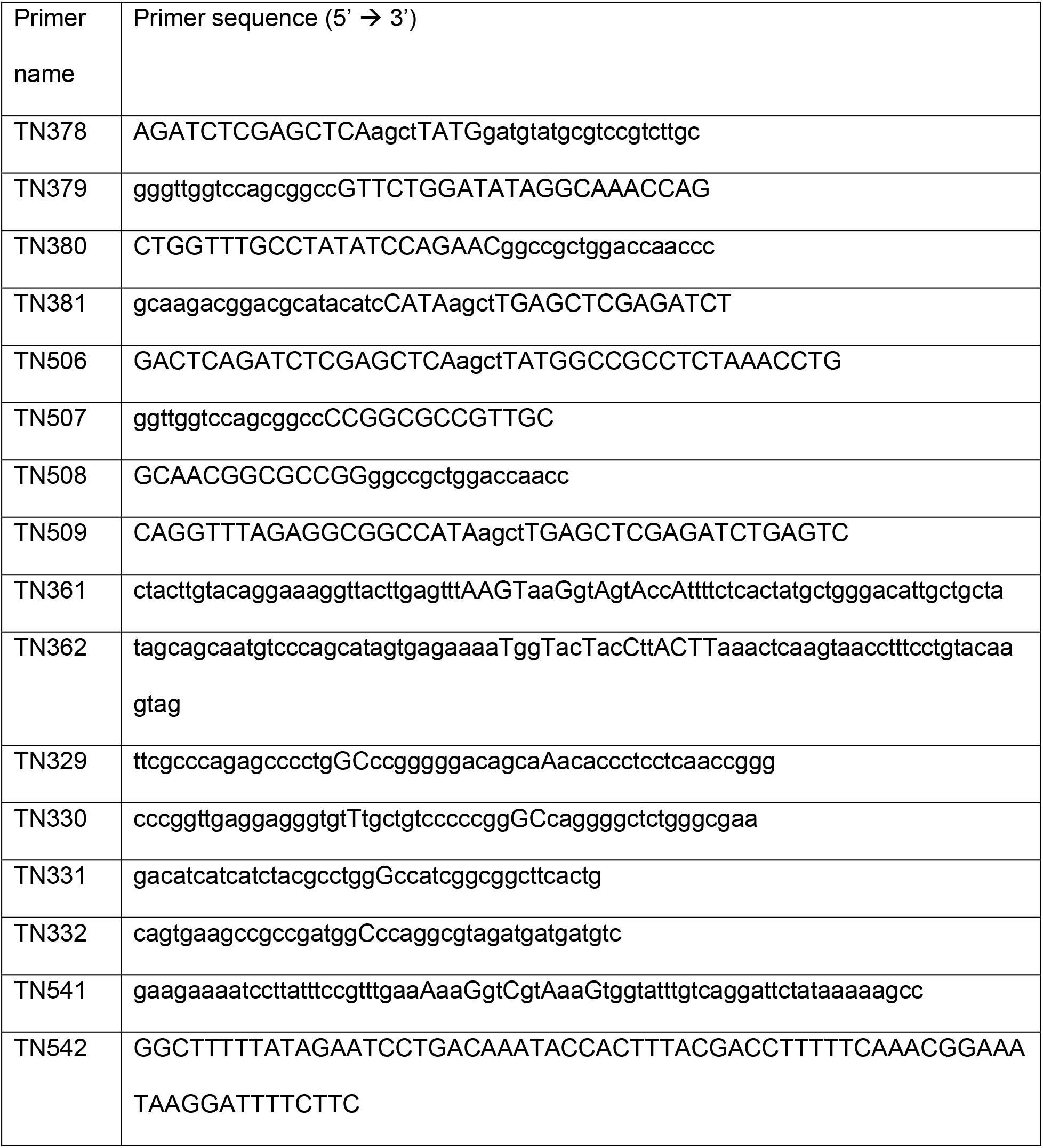

### General reagents

Mouse anti-V5 tag monoclonal antibody (Thermo Fisher Scientific, Cat# R960-25, RRID: AB_2556564) was used at 1:2000 for western blot and 1:200 for IF.

Rabbit anti-calnexin polyclonal antibody (Enzo Life Sciences, Cat# ADI-SPA-860-F, RRID: AB_11178981) was used at 1:4000 for western blot.

Rabbit anti-calreticulin polyclonal antibody (Abcam, Cat# ab2907, RRID: AB_303402) was used at 1:2000 for western blot.

Rabbit anti-GAPDH antibody (Sigma-Aldrich, Cat# G9545-200UL, RRID: AB_796208) was used at 1:100,000 for western blot.

Mouse anti-tom20 antibody (F-10) (Santa Cruz Biotechnology, Cat# sc-17764, RRID: AB_628381) was used at 1:500 for western blot.

Rabbit anti-COX IV monoclonal antibody (Cell Signaling Technology, Cat# 4850S) was used at 1:1000 for western blot.

Rabbit anti-ABHD16A antibody (Abcam, Cat# ab185549) was used at 1:1000 for western blot. Rabbit anti-ORP8 polyclonal antibody (GeneTex, Cat# GTX121273) was used at 1:1000 for western blot.

Rabbit anti-VPS13A (Chorein) polyclonal antibody (Novus Biologicals, Cat# NBP1-85641) was used at 1:1000 for western blot.

Rabbit anti-VPS13D polyclonal antibody (Abcam, Cat# ab202285) was used at 1:1000 for western blot.

Donkey anti-Mouse IgG (H+L) Highly Cross-Adsorbed Secondary Antibody, Alexa Fluor 647 (Thermo Fisher Scientific, Cat# A-31571, RRID:AB_162542) was used at 1:200 for IF.

Goat anti-rabbit IgG antibody, HRP-conjugate (Sigma-Aldrich, Cat. #12-348) was used at 1:6000 for western blot.

Goat anti-mouse IgG antibody, HRP-conjugate (Sigma-Aldrich, Cat. # 12-349) was used at 1:3000 for western blot.

4-20% Criterion TGX Precast Midi Protein Gels (Bio-Rad, Cat. #5671094 and #5671024) were used to run western blots.

SuperSignal West Pico PLUS Chemiluminescent Substrate (Thermo Fisher, Cat. # 34577) was used to develop western blots.

Standard SDS-PAGE/Western protocols were used to develop western blots.

### Cell culture and transfections

U-2 OS cells (ATCC) were cultured in McCoy’s 5A (Invitrogen, Cat# 16600-108) with 10% FBS (Sigma-Aldrich, Cat# 12306C-500ML) and 1% Penicillin-Streptomycin (Invitrogen, Cat# 15070063) at 37°C with 5% CO_2_. HeLa cells (ATCC) were cultured in DMEM (Gibco, Cat# 12430-062) with 10% FBS (Sigma-Aldrich, Cat# 12306C-500ML) and 1% Penicillin-Streptomycin (Invitrogen, Cat# 15070063) at 37°C with 5% CO_2_.

For imaging, cells were plated on 35 mm imaging dishes (Cellvis, Cat. #D35-10-1.5-N) for 16-20 hours, then transfected for 5 hours with indicated plasmids in 2 mL Opti-MEM (Invitrogen, Cat. #31985-088) using Lipofectamine 3000 transfection kit (Thermo Fisher, Cat. #L3000-150). Cells were imaged 16-20 hrs after transfection in FluoroBrite DMEM (Gibco, Cat. #A18967-01) supplemented with 10% FBS, 1% Penicillin-Streptomycin, 1% GlutaMAX (Gibco, Cat. #35050-061), and 25 mM HEPES. If imaging SNAP-tag or Halo tag, cells were incubated with Janelia Fluor 646, SE (1.5 μM in DMSO) in serum-free FluoroBrite DMEM with added supplements for 30 minutes prior to imaging or Janelia Fluor X 646 in complete FluoroBrite DMEM with added supplements for 30 minutes prior to imaging. Janelia Fluor 646, SE and Janelia Fluor X 646 were gifts from Luke Lavis, Janelia Research Facility.

For transfection, plasmids were incubated at room temperature with 2 μL P3000 per μg of plasmid in 250 μL Opti-MEM while 2.5 μL/mL (of Opti-MEM) Lipofectamine 3000 was also incubated in 250 μL Opti-MEM. After 5 minutes, plasmids + P3000 and Lipofectamine 3000 were mixed together and incubated at room temperature for 20 minutes. Mixture was added drop-wise onto cells in 1.5 mL Opti-MEM for 5 hours.

For plasmid concentrations, we used 125 ng/mL (for IF) 250 ng/mL (for mass spectrometry) V5-TurboID-Mfn1; 150 ng/mL (for IF) 50 ng/mL (for mass spectrometry) V5-TurboID-Mfn1E209A; 125 ng/mL GFP-Mfn1; 75 ng/mL mito-BFP, 25 ng/mL (low expression) 250 ng/mL (overexpression) Aphyd-mNG; 75 ng/mL (KD+rescue) 250 ng/mL (overexpression) mNG-N1; 200 ng/mL mCherry-Sec61β; 75 ng/mL (KD+rescue) 250 ng/mL (overexpression) Stim1NTD-Aphydcyto-mNG (Aphyd^ER^); 75 ng/mL (KD+rescue) 250 ng/mL (overexpression) Tom70NTD-Aphydcyto-mNG (Aphyd^mito^); 75 ng/mL siRNA-resistant Aphyd-mNG; 100 ng/mL mCherry-N1; 100 ng/mL or 250ng/mL (overexpression) siRNA-resistant Aphyd-mCherry; 200 ng/mL mNG-Sec61β; 125 ng/mL mNG-Mfn1; 75 ng/mL Halo-N1; 75 ng/mL siRNA-resistant Aphyd-Halo; 150 ng/mL mMAPLE-OMP25; 125 ng/mL RA-Sec61β; 125 ng/mL B-Mff; 75 ng/mL SNAP-N1; 75 ng/mL siRNA-resistant Aphyd-SNAP; 75 ng/mL siRNA-resistant Aphyd-mNG acyltransferase motif mutant (H189AD194N); 75 ng/mL siRNA-resistant Aphyd-mNG alpha/beta hydrolase domain mutant (S355A); 75 ng/mL siRNA-resistant Aphyd-Halo acyltransferase motif mutant (H189AD194N); 75 ng/mL siRNA-resistant Aphyd-Halo alpha/beta hydrolase domain mutant (S355A); 125 ng/mL siRNA-resistant mNG-ORP8.

For KD +/−rescue experiments, cells were treated with siRNAs twice. The following siRNA oligonucleotides were used: Silencer Negative Control #1 siRNA (Ambion, Cat. #AM4635) used at 25 nM and Aphyd siRNA (Horizon Discovery, Cat# J-013106-09-0010) used at 25 nM against target sequence: 5’-UGUCCAAAGUGGUGCCGUU-3’. The Aphyd siRNA used targets a region inside the ORF so all Aphyd constructs were mutated to be resistant. OSBPL8 siRNA (Horizon Discovery, Cat# J-009508-07-0005) used at 25 nM against target sequence: 5’-AGAAAGUAGUGAAAUGGUA-3’. The OSBPL8 siRNA used targets a region inside the ORF so siRNA-resistant mNG-ORP8 was mutated to be resistant. VPS13A SMARTpool siRNA (Horizon Discovery, Cat# L-012878-00-0005) used at 25 nM against target sequences: 5’-GGAUAGAGCUUAUGAUUCA-3’, 5’-GAAUGGCACUGGAUAUUAA-3’, 5’-UAACACAUCUGCACAUCAA-3’, 5’-GCAGCUACAUUCCUCUUAA-3’. VPS13D SMARTpool siRNA (Horizon Discovery, Cat# L-021567-02-0005) used at 25 nM against target sequences: 5’-UCUAAGAACUGCCGAGAAU-3’, 5’-CAAGAAAGGCCGAGGUCGA-3’, 5’-GGAAGGCAGUGCACGGAAA-3’, 5’-AUGUUAAGACUCAGCGAAA-3’. First, cells were plated in 6-well plates (Greiner Bio-One Cellstar, Cat. #657-160) for 16-20 hours then transfected with indicated siRNA oligonucleotides with DharmaFECT 1 Transfection Reagent (Cat. #T-2001-02) for the first round of siRNA treatment. Indicated concentrations of siRNA were incubated in 250 μL serum-free, antibiotic-free media while 5 μL DharmaFECT 1 Transfection Reagent was incubated in 250μL serum-free, antibiotic-free media separately at room temperature. After 5 minutes, siRNA and DharmaFECT 1 mixtures were mixed together and incubated at room temperature for 20 minutes. Mixture was added drop-wise onto cells into 1.5mL serum-free antibiotic-free media for 6 hrs. The following day, cells were trypsinized with 0.25% Trypsin-EDTA (Gibco, Cat. #25200-072) and plated into 6-well dishes for immuno-blot analysis or 35mm imaging dishes for 16-20 hrs. The second round of transfection was done in Opti-MEM with the same concentration of siRNA oligonucleotides and plasmids necessary for imaging using only Lipofectamine 3000 without P3000 for 5 hrs. Cell lysate was collected and images were taken 16-20 hrs after transfection. Cell lysate was collected by directly lysing in 2x Laemmli Sample Buffer (BioRad, Cat. #1610737) and boiling for 10 minutes at 95°C after trypsinizing and washing once with 1X Dulbecco’s Phosphate Buffered Saline (Sigma Aldrich, Cat. #D1408-500ML).

### Immunofluorescence

HeLa cells were seeded on 35 mm imaging dishes (Cellvis, Cat. #D35-10-1.5-N). 16-20 hours after seeding, cells were transfected with V5-TurboID-Mfn1 or V5-TurboID-Mfn1E209A, GFP-Mfn1, and mito-BFP. 16-20 hours after transfection, cells were washing once with 1x PBS and fixed with 4% paraformaldehyde in 1x PBS for 15 minutes. Cells were permeabilized with 0.1% TX100 in 1x PBS for 5 minutes. Cells were washed 3 times with 1x PBS and blocked in 10% normal donkey serum and 0.1% TX100 in 1x PBS (blocking buffer) for 60 minutes. Primary antibody against the V5 tag in blocking buffer was added over night. The next day, cells were washed 3 times with 1x PBS and stained with secondary antibody in blocking buffer for 30 minutes. Cells were washed 3 times with 1x PBS and imaged on the spinning disk confocal.

### Microscopy

All cells were imaged on a spinning disk confocal microscope except for Figure 3, which was on a Zeiss LSM 880 Confocal Laser Scanning Microscope equipped with a Plan-Apochromat 63x oil objective (1.4 N.A) and Airyscan detector and controlled by Zen Black (Zeiss). The spinning disk is a Nikon Ti2E body with PSF; CSU-X1 Yokogawa confocal scanner unit; OBIS 405, 488, 561, 640 lasers and an ANDOR iXON Ultra 512×512 EMCCD camera. Images were acquired with Nikon Plan Apo λ 100X oil objective (1.4 N.A). Images were acquired with micro-manager 2.0. Images were processed in Fiji (NIH) and Adobe Illustrator (Adobe).

### Image collection and quantification

For mitochondrial area quantification, z-stacks consisting of 12 serial images that were each spaced by 0.2 μm. On Fiji, 15×15 μm ROIs were drawn around resolvable regions of mitochondria to assess morphology. Then, mitochondria were individually counted using the ‘Multi-point’ tool. The total area of the mitochondria in the ROI was calculated by Otsu Thresholding. Thresholded images were then assessed using the ‘Analyze Particles’ function to obtain the total area in the ROI. Finally, the total mitochondrial area was divided by the number of mitochondria in the ROI to obtain the average area per mitochondrion (μm^2^).

For ER crossings containing Drp1 puncta, 2-minute movies were taken at 5 second intervals. Resolvable ER-mitochondria crossings were marked with the ‘Multi-point’ tool on Fiji. Presence of a Drp1 puncta were counted (as assessed by puncta present throughout a 2-minute movie) and the total number of Drp1 puncta was divided by the total number resolvable ER-mitochondria crossings to get the percentage of ER crossings containing Drp1 puncta.

For Mfn1 density quantifications, 3 lines were drawn along the length of resolvable mitochondrion per cell using the ‘Segmented line’ tool in Fiji to obtain mitochondrial length. Then, Mfn1 puncta were counted (as assessed by puncta present throughout a 2-minute movie), and the total number of Mfn1 puncta along one mitochondrion was divided by length of the one mitochondrion to give number of Mfn1 puncta per micron (Mfn1 density).

For assessing fission and fusion rate, cells were transfected with mMAPLE-OMP25 as described. Single mitochondria were photoconverted from green to red fluorescence by stimulating with 20 iterations of 100% 405 nm light on the Zeiss LSM 880 Confocal Laser Scanning Microscope. Apparent fusion/fission was scored during live 5-minute time-lapse movies with 5 second intervals by the observation of fluorescence mixing or fluorescence separation. The fusion/fission rate was calculated by dividing the number of fusion/fission events per mitochondrion per minute. Mitochondrial fusion was binned in two categories: tip-to-tip and tip-to-middle fusion within these experiments.

For mitochondria overlap with ER or MCSs, 20×20 μm or 15×15 μm ROIs were selected for resolvable regions of ER, mitochondria, and ddFP signal. Then, the ‘JaCoP’ plugin for Fiji was used to manually threshold each image and calculate Mander’s correlation coefficient (MCC) for the percentage of ER covering mitochondria signal or mitochondria covering ddFP signal.

For mitochondrial constriction line scans and quantifications, z-stacks consisting of 12 serial images that were each spaced by 0.2 μm were taken. Resolvable ER-mitochondria crossings were marked with the ‘Multi-point’ tool on Fiji. Line scans were performed using Fiji by drawing a line along the mitochondrion and the fluorescence along the mitochondrion was measured. Constrictions were marked as a ≥40% decrease in mitochondrial signal intensity compared to the neighboring fluorescence peak. The number of mitochondrial constrictions was divided by the total number of resolvable ER-mitochondria crossings to get the percentage ER crossings containing constrictions.

### Statistics

All data presented in all figures are from at least three biological replications. All figure quantifications are represented as violin plots with the bolded line representing the median and the peripheral lines representing quartiles. All statistical analyses were performed in GraphPad Prism 8. When comparing two samples, two-tailed Student’s t tests were used. When analyzing significance for more than two samples, one-way ANOVA tests were performed and p values were derived from Tukey’s test. ns = not significant, **p < 0.01, ***p < 0.001, ****p < 0.0001. Details of statistical analysis and exact values of n (numbers of cells) quantified in each experiment can be found in the figure legends.

### Biotinylation, ER isolation, and biotinylated protein collection

HeLa cells plated in 20 10-cm dishes ~16-20 hours prior to transfection to attain ~90% confluency the day of biotinylation and sample collection. Cells were transfected with V5-TurboID-Mfn1 or V5-TurboID-Mfn1E209A one day after plating. ~16 hours after transfection, cells were treated with 500 μM biotin for 3 hours. After biotin treatment, cells were washed once with cold 1x PBS, trypsinized, and pelleted. Cell pellets were washed twice with cold 1x PBS and resuspended in 2m L IB-1 with cOmplete™, Mini, EDTA-free Protease Inhibitor Cocktail (Sigma-Aldrich, Cat# 11836170001) (225 mM mannitol, 75 mM sucrose, 30 mM Tris-HCl pH=7.4, 0.1 mM EGTA) (Wieckowski et al., 2009). Cells were lysed via sonication for 10 seconds 3 times. Lysed cells were spun at 600 x g for 5 minutes twice to rid of whole cells, debris, and the nuclear fraction. The supernatant was then spun at 7000 x g for 20 minutes twice to rid of the mitochondrial fraction. The supernatant was then spun at 20,000 x g for 30 minutes. The pellet (containing the ER fraction) after the 20,000 x g spin was resuspended in IB-2 (225 mM mannitol, 75 mM sucrose, 30 mM Tris-HCl pH=7.4) and spun again at 20,000 x g for 30 minutes to rid of other contaminating fractions. The supernatant was taken and spun at 100,000 x g for 30 minutes to pellet the residual ER fraction. The pellet (containing the ER fraction) after the 100,000 x g spin was resuspended in IB-2 (225 mM mannitol, 75 mM sucrose, 30 mM Tris-HCl pH=7.4) and spun again at 100,000 x g for 30 minutes to rid of other contaminating fractions. The cleaned 20,000 and 100,000 x g pellets were resuspended in MRB (225 mM mannitol, 75 mM sucrose, 30 mM Tris-HCl pH=7.4, 0.1 mM EGTA) and 0.1% SDS, flash frozen in liquid nitrogen, and sent for mass spectrometry.

### Sample preparation for mass spectrometry

All sample preparation and mass spectrometry were conducted at the proteomics core at Sanford Burnham Prebys. Protein concentration was determined using a bicinchoninic acid (BCA) protein assay (Thermo Scientific). Disulfide bridges were reduced with 5 mM tris (2-carboxyethyl)phosphine (TCEP) at 30°C for 60 min, and cysteines were subsequently alkylated with 15 mM iodoacetamide (IAA) in the dark at room temperature for 30 min. Affinity purification was carried out in a Bravo AssayMap platform (Agilent) using AssayMap streptavidin cartridges (Agilent). Briefly, cartridges were first primed with 50 mM ammonium bicarbonate, and then proteins were slowly loaded onto the streptavidin cartridge. Background contamination was removed with 8 M urea, 50 mM ammonium bicarbonate. Finally, cartridges were washed with Rapid digestion buffer (Promega, Rapid digestion buffer kit) and proteins were subjected to on-cartridge digestion with mass spec grade Trypsin/Lys-C Rapid digestion enzyme (Promega, Madison, WI) at 70°C for 1 hour. Digested peptides were then desalted in the Bravo platform using AssayMap C18 cartridges and dried down in a SpeedVac concentrator.

### LC-MS/MS

Prior to LC-MS/MS analysis, dried peptides were reconstituted with 2% ACN, 0.1% FA and concentration was determined using a NanoDropTM spectrophometer (ThermoFisher). Samples were then analyzed by LC-MS/MS using a Proxeon EASY-nanoLC system (ThermoFisher) coupled to a Q-Exactive Plus mass spectrometer (Thermo Fisher Scientific). Peptides were separated using an analytical C18 Aurora column (75μm x 250 mm, 1.6 μm particles; IonOpticks) at a flow rate of 300 nL/min (60C) using a 120-min gradient: 1% to 5% B in 1 min, 6% to 23% B in 72 min, 23% to 34% B in 45 min, and 34% to 48% B in 2 min (A= FA 0.1%; B=80% ACN: 0.1% FA). The mass spectrometer was operated in positive data-dependent acquisition mode. MS1 spectra were measured in the Orbitrap in a mass-to-charge (m/z) of 350 – 1700 with a resolution of 70,000 at m/z 400. Automatic gain control target was set to 1 x 106 with a maximum injection time of 100 ms. Up to 12 MS2 spectra per duty cycle were triggered, fragmented by HCD, and acquired with a resolution of 17,500 and an AGC target of 5 x 104, an isolation window of 1.6 m/z and a normalized collision energy of 25. The dynamic exclusion was set to 20 seconds with a 10 ppm mass tolerance around the precursor.

### Mass spectrometry data analysis

All mass spectra from were analyzed with MaxQuant software version 1.5.5.1. MS/MS spectra were searched against the Homo sapiens Uniprot protein sequence database (downloaded in January 2020) and GPM cRAP sequences (commonly known protein contaminants). Precursor mass tolerance was set to 20ppm and 4.5ppm for the first search where initial mass recalibration was completed and for the main search, respectively. Product ions were searched with a mass tolerance 0.5 Da. The maximum precursor ion charge state used for searching was 7. Carbamidomethylation of cysteine was searched as a fixed modification, while oxidation of methionine and acetylation of protein N-terminal were searched as variable modifications. Enzyme was set to trypsin in a specific mode and a maximum of two missed cleavages was allowed for searching. The target-decoy-based false discovery rate (FDR) filter for spectrum and protein identification was set to 1%.

